# Spatio-temporal modeling of high-throughput multi-spectral aerial images improves agronomic trait genomic prediction in hybrid maize

**DOI:** 10.1101/2022.10.18.512728

**Authors:** Nicolas Morales, Mahlet T. Anche, Nicholas S. Kaczmar, Nicholas Lepak, Pengzun Ni, Maria Cinta Romay, Nicholas Santantonio, Edward S. Buckler, Michael A. Gore, Lukas A. Mueller, Kelly R. Robbins

## Abstract

Design randomizations and spatial corrections have increased understanding of genotypic, spatial, and residual effects in field experiments, but precisely measuring spatial heterogeneity in the field remains a challenge. To this end, our study evaluated approaches to improve spatial modeling using high-throughput phenotypes (HTP) via unoccupied aerial vehicle (UAV) imagery. The normalized difference vegetation index (NDVI) was measured by a multi-spectral MicaSense camera and ImageBreed. Contrasting to baseline agronomic trait spatial correction and a baseline multi-trait model, a two-stage approach that quantified NDVI local environmental effects (NLEE) was proposed. Firstly, NLEE were separated from additive genetic effects over the growing season using two-dimensional spline (2DSpl), separable autoregressive (AR1) models, or random regression models (RR). Secondly, the NLEE were leveraged within agronomic trait genomic best linear unbiased prediction (GBLUP) either modeling an empirical covariance for random effects, or by modeling fixed effects as an average of NLEE across time or split among three growth phases. Modeling approaches were tested using simulation data and Genomes-to-Fields (G2F) hybrid maize (*Zea mays* L.) field experiments in 2015, 2017, 2019, and 2020 for grain yield, grain moisture, and ear height. The two-stage approach improved heritability, model fit, and genotypic effect estimation compared to all baseline models. Electrical conductance and elevation from a 2019 soil survey significantly improved model fit, while 2DSpl NLEE were most correlated to the soil parameters and grain yield 2DSpl effects. Simulation of field effects demonstrated improved specificity for RR models. In summary, NLEE increased experimental accuracy and understanding of field spatio-temporal heterogeneity.

## Introduction

The importance of controlling for environmental heterogeneity in agricultural field experiments is critical to obtain accurate estimates of varietal performance and treatment effects (Smith, Cullis, and Thompson 2005; Van Es and Van Es 1993; Brownie, Bowman, and Burton 1993; Xu 2016). In plant breeding where soil composition, elevation, slope, curvature, water content, nutrient availability, and management can vary within field experiments, the genotypic effects driving important agronomic traits become confounded with local environment effects (LEE). Randomization in experimental designs can help control errors from spatial variation to a large degree (Piepho, Möhring, and Williams 2013; Hoefler et al. 2020), but in early stage trials where replication is limited, it is important to account for spatial variation. Statistical approaches, such as the separable autoregressive process and the two-dimensional spline model, have advanced to capture local dependence effects between experimental plots (Arthur R. Gilmour et al. 1997; Covarrubias-Pazaran 2016). Such spatial effects derive local dependencies from distance-based random covariance structures (e.g. plots that are close to each other are more interdependent than those farther away), but these models often make simplifying assumptions of a consistent rate of decay in interdependency across the entire field. Nonetheless, statistically modeling spatial effects using linear mixed models has improved experimental accuracy in plant breeding (Rodríguez-Álvarez et al. 2018; Robbins, Backlund, and Schnelle 2012; Copati et al. 2021; Bernardeli et al. 2021; Smith, Cullis, and Thompson 2005).

Spatial heterogeneity can change over the growing season, due to weather and management conditions as well as plant development characteristics. The relationship of time on spatial effects can be explored through univariate and multivariate spatial models. Also, repeated measurements in time allow estimation of permanent environment (PE) effects from random regression (RR) models, providing a purely temporal representation of spatial heterogeneity. To effectively apply these statistical approaches, measurements should largely span the field; however, it can be difficult and expensive to objectively measure phenotypic traits repeatedly across a field experiment, as seen with quantitative disease resistance traits (Poland and Nelson 2011; Reynolds et al. 2019). Therefore, cost-effective remote sensing approaches are necessary for studying LEE in the field across time.

Through the use of unoccupied aerial vehicles (UAVs) and other systems, aerial imaging can reliably and cost-effectively measure high-throughput phenotypes (HTPs) for all experimental plots in the field across the growing season (White et al. 2012; Andrade-Sanchez et al. 2014; Sagan et al. 2019; Sun et al. 2021). A widely studied class of aerial image HTPs are vegetation indices (VI) that include the normalized difference vegetation index (NDVI) (Gitelson et al. 2002; Hunt et al. 2013). VI provide physiologically relevant image features that can track variance such as photosynthetic activity, and have successfully measured chlorophyll content, canopy extent, biomass, and water use efficiency among other plant attributes (Thorp et al. 2018; Delegido et al. 2011; Bannari et al. 2007; Babar et al. 2006). Promising model-derived HTPs from images exist, such as latent-space and convolutional neural network (CNN) features; however, our study focused on NDVI LEE (NLEE) from linear mixed models to offer a standardized approach for quantifying latent spatial heterogeneity and field environmental effects (Gage, Richards, and Lepak 2019; Taghavi Namin et al. 2018; Wiesner-Hanks et al. 2019; Feldmann et al. 2021).

While phenotypic data are critical in plant breeding, genomic data are arguably of equal importance. Applying whole-genome marker data for genomic prediction (GP) is now feasible with the proliferation of genotyping technologies (Meuwissen, Hayes, and Goddard 2001). Genomic best linear unbiased prediction (GBLUP) has been extensively applied to predict traits in animals and plants from genome-wide single nucleotide polymorphism (SNP) markers, including in maize and wheat (Rutkoski et al. 2012; Daetwyler et al. 2013). GBLUP can predict traits from genome-wide SNP marker data by modeling the covariance of additive genetic effects as a genomic relationship matrix (GRM) (VanRaden 2008).

VI can improve GP through multivariate approaches by leveraging genetic correlations between the VI and agronomic traits of interest, as demonstrated for grain yield in wheat and biomass in soybean (Rutkoski et al. 2016; Sakurai et al. 2021). While multi-trait models leverage genetic correlations across traits to improve predictions, residual correlations exist between NDVI and grain yield in maize (Anche et al. 2020). Recent studies have successfully proposed two-stage approaches for incorporating HTP, such as detecting spatial effects using the SpATS package in the first stage and then creating P-spline hierarchical growth models in the second stage (Pérez-Valencia et al. 2022). Furthermore, modeling approaches for integrating HTP, genomic information, and environmental information can be generalized into the genotype-to-phenotype (G2P) model framework (van Eeuwijk et al. 2019); however, modeling NLEE into GP has not previously been widely explored and field tested.

Building on previous work, our study proposed a two-stage approach for improving spatial corrections in GP. The first stage separated NLEE from additive genetic effects in the HTP temporally, using either spatial corrections or RR models, to quantify the LEE in the field. The second stage summarized the NLEE within GBLUP for the agronomic traits using two distinct implementations, either modeling a plot-to-plot covariance of random effects (L) or modeling fixed effects (FE). The proposed approach studied the following two questions utilizing simulated data and several years of hybrid maize field experiments. Firstly, are NLEE, estimated by spatial effects and PE effects, consistently able to detect across the growing season the spatial heterogeneity affecting end-of-season agronomic traits? Secondly, can NLEE be used in the proposed two-stage models to improve spatial corrections for GP of agronomic traits?

## Materials and Methods

### Field Experiments

As part of the Genomes to Fields (G2F) program, inbred and hybrid maize (*Zea mays* L.) field evaluations were planted at the Musgrave Research Farm (MRF) in Aurora, NY (McFarland et al. 2020). Of importance to this study were the hybrid maize experiments planted in 2015 (https://doi.org/10.25739/erxg-yn49), 2017 (https://doi.org/10.25739/w560-2114), 2019 (https://doi.org/10.25739/t651-yy97), and 2020 (https://doi.org/10.25739/hzzs-a865), named 2015_NYH2, 2017_NYH2, 2019_NYH2, and 2020_NYH2, respectively. Each experimental plot was seeded in two-row plantings. In all field experiments, the following agronomic traits were measured: grain yield (GY) (bu/acre), grain moisture (GM) (%), and ear height (EH) (cm).

Experimental variation across the years arose from the maize hybrids planted, the field sites utilized, the weather conditions experienced during the growing seasons, and the time points at which imaging events occurred. Figure S1 illustrated the time points at which high-throughput phenotypes (HTP) were extracted, as well as the growing degree days (GDD) and cumulative precipitation (CP) on those days. GDD were calculated for each imaging event time point from the planting date of the experiment and were found using daily weather data for the ground station at MRF (GHCND:USC00300331) accessed via the NOAA NCEI NCDC database. The evidenced variation in GDD and CP across years highlighted the necessity to collect as many imaging events as possible over the growing season.

### Aerial Image Collection and Processing

A MicaSense RedEdge 5-channel multi-spectral camera mounted onto an unoccupied aerial vehicle (UAV) captured images in the blue, green, red, near infrared, and red-edge spectra. The UAV flew at an altitude of 25m to 30m and at a speed of 6 km/hr. To complete a flight, the pre-programmed, serpentine flight plans required approximately 35 min to traverse the 3km path. At least 80% overlap along both image axes was ensured in the collected images.

Each flight produced approximately 5,000 images from the MicaSense camera. The images were then processed into orthophotomosaics using Pix4dMapper photogrammetry software. To produce reflectance calibrated raster images, this software used the MicaSense radiometric calibration panel images captured immediately prior to each UAV flight, as well as illumination metadata embedded in each capture by the MicaSense camera. Orthophotomosaic images were produced with approximately 1cm per pixel resolution ground sample distance (GSD). The resulting reflectance orthophoto images were then uploaded into ImageBreed, which enabled plot-polygon templates to be created and assigned to the field experimental design (Morales, Kaczmar, et al. 2020). Figure S2 illustrates a representative near-infrared (NIR) reflectance orthophoto image from 2019_NYH2 taken on August 15, 2019, with the plot-polygons overlaid. Normalized difference vegetation index (NDVI) HTP were extracted from ImageBreed, derived from the plot image mean pixel value (Gitelson et al. 2002; Hunt et al. 2013; Patrignani and Ochsner 2015; Bhandari et al. 2021). The image, field experiment, phenotypic, and genotypic data within ImageBreed are FAIR and queryable through openly described APIs (Selby et al. 2019; Wilkinson et al. 2016).

Flights in 2015, 2017, and 2019 were scheduled approximately once per week, while 2020 targeted a frequency of twice per week. Technical problems and poor weather conditions, such as clouds, rain, and high winds, resulted in fewer imaging events being suitable for HTP extraction. To separate the HTP imaging event dates into biological growth stages, GDD ranges were defined to account for the early vegetative phase (P0) at 0 to 1225 GDD, the active reproductive phase (P1) at 1226 to 1800 GDD, and the late reproductive phase (P2) at 1801 to 2500 GDD. The three GDD ranges captured periods where NDVI was first steadily increasing, then plateauing, and finally steadily decreasing. Table 1 summarized the growth stage distribution of imaging event dates for which HTP were successfully extracted.

**Table 1:** Summary of UAV imaging dates for which HTP were successfully extracted in the 2015, 2017, 2019, and 2020 field experiments. Growth stages of P0, P1, and P2 map to early, active, and late reproductive phases broadly defined as 0 to 1225 GDD, 1226 to 1800 GDD, and 1801 to 2500 GDD, respectively.

| Field Experiment | Planting Date | Field Location | Growth Stage: P0 | Growth Stage: P1 | Growth Stage: P2 |
| --- | --- | --- | --- | --- | --- |
| 2015_NYH2 | May 7, 2015 | MRF, Field P | July 21 | Aug 7, Aug 20 | Sept 10 |
| 2017_NYH2 | May 18, 2017 | MRF, Field Y | June 12 | Aug 2, Aug 17, Sept 1 | Sept 6, Sept 12, Sept 24 |
| 2019_NYH2 | May 23, 2019 | MRF, Field N | July 16, July 24, July 29 | Aug 5, Aug 15 | Sept 10 |
| 2020_NYH2 | May 22, 2020 | MRF, Field D | June 29, July 9, July 15, July 18 | July 22, July 28, Aug 1 | Aug 20, Aug 26, Sept 9, Sept 18, Oct 2 |

### Soil Information

In 2019 at the MRF, a ground conductivity meter (EM38-MK2, Geonics, Canada) surveyed Field N where the G2F hybrid maize field experiment was planted. The EM-38 probe used electrical inductance to characterize variation originating from a combination of factors including soil salinity, soil texture, water content, water retention, soil type, and soil nutrients (Heil and Schmidhalter 2017). The goals for the soil information in this study were to (1) better understand driving factors for the detected NLEE from the aerial imagery, and (2) determine whether a soil survey was a practical alternative to aerial imagery for improving agronomic GP in the second stage. Due to planting rotations and other logistical concerns, the G2F experiments conducted in 2015, 2017, 2019, and 2020 were all in distinct field locations as shown in Table 1; therefore, the soil information in this study could only be applied to the 2019 experiment.

The soil survey was conducted prior to the hybrids being planted by passing the probe over the field in a dual-serpentine pattern, with 9 passes in the east-west orientation and 24 passes in the north-south orientation. The georeferenced elevation (Alt) and apparent electrical conductance (EC) data were then interpolated over the entire field using ordinary Kriging in R (Pebesma 2004). The interpolated raster was produced using a spherical variogram model at a resolution of 10^-6^ WGS84 units covering 120 by 200 cells. Figure S3 illustrates (A) a map of the collected EM38 soil survey, (B) the region to interpolate into, and in (C) and (D) the interpolated soil EC and Alt across the field, respectively. Finally, a mean value for the soil EC and Alt was extracted using ImageBreed for each plot in the 2019_NYH2 field experiment.

In order to approximate soil curvature as elevation gradients, first and second two-dimensional numerical derivatives were computed on the plot-level soil EC and altitude measurements, denoted as *dEC* (dEC), *d*^2^*EC* (d2EC), *dAlt* (dAlt), and *d* ^2^*Alt* (d2Alt), respectively. Two-dimensional numerical derivatives were computed by averaging the differences between a given plot and the three immediately adjacent rings encircling it. Figure S4 illustrated heatmaps of the extracted plot-level soil EC and Alt, along with the first and second derivatives.

### Genotyping Data

Through the G2F program, genotype-by-sequencing (GBS) resulted in 945,574 SNP markers across the genome for 1577 samples representing a total of 1325 unique maize inbred lines (https://doi.org/10.25739/frmv-wj25) (McFarland et al. 2020; Elshire et al. 2011). The resulting genome-wide variant call format (VCF) data were queried for the hybrids in the 2015, 2017, 2019, and 2020 field experiments (Morales, Bauchet, et al. 2020; Morales et al. 2022; Danecek et al. 2011). Due to minor typographical errors (e.g. Mo17 vs MO17), data cleaning was required prior to mapping the sample identifiers in the VCF to the field experiment genotype identifiers and to the pedigree information for the maize hybrids. Genotypes were filtered for SNPs with minor allele frequency < 5% or with > 40% missing data, and for samples containing > 20% missing data. Genomic relationship matrices (GRMs) were computed using the A.mat function in rrBLUP, with missing data imputed as the mean genotype (Endelman 2011; VanRaden 2008); therefore, only an additive genetic relationships were modeled (Griffing 1956).

Given that many of the evaluated maize hybrids in the G2F program originated as bi-parental crosses of the genotyped inbred lines and that the inbred lines are unrelated to each other, hybrid maize genotypes were computed by averaging over the parental GRMs. Specifically, the GRMs among the pollen parents and seed parents were computed independently, and then the hybrid’s relationship was computed as an average among one half of the inbred parental relationships. If a hybrid’s parental inbred lines were not genotyped, then the hybrid was included in the GRM with a diagonal value of one and off-diagonal values of zero.

### HTP Spatial Heterogeneity

The first question of this study was whether NDVI LEE (NLEE), which was estimated by spatial and PE effects, consistently detected across the growing season the spatial heterogeneity affecting end-of-season agronomic traits. Therefore, the first stage of the proposed two-stage approach focused on measuring spatial heterogeneity in NDVI across the growing season.

Variance in the collected HTP observations,*V*_*P*_, was modeled as arising from genetic variance among the maize hybrids,*V*_*G*_, and from environmental sources of variance,*V*_*E*_ (Falconer and Mackay 2009). In mixed model matrix notation this was formulated as

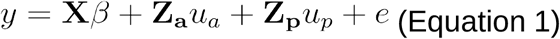

where *y* was a vector of phenotypic observations and X was an incidence matrix mapping phenotypic values to the fixed effects, *β*. The random effects *u*_*a*_ and *u*_*p*_ represented the additive genetic and local environment, respectively, and the incidence matrices Z_a_ and Z_p_ linked the random effects *u*_*a*_ and *u*_*p*_ respectively, to the observations.

The spatial model fitted either a single HTP time point or multiple HTP time points, a distinction referred to as the univariate or multivariate cases. In the univariate case, spatial models were run independently for each of the collected HTP time points. In contrast, the multivariate case fitted several collected HTP time points in a single model, such that, the vector *y* represented HTP observations taken at different time

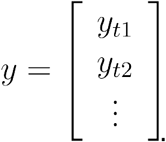

points across the growing season,

In the univariate spatial case, the variance of the random additive genetic effect was defined as

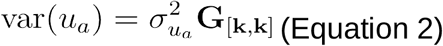

where the matrix G represented the GRM between evaluated hybrids and was of order [*k,k*] denoting the total number of hybrids *k* (VanRaden 2008). The multivariate case defined this as

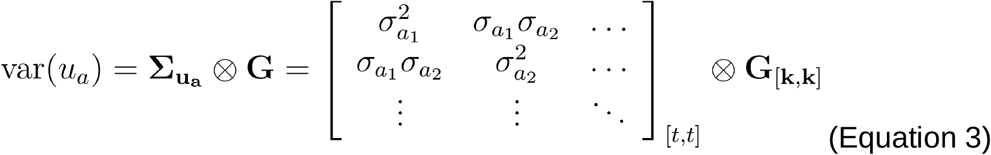

where the unstructured matrix 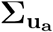 was of order [*t,t*]denoting the number of time points involved and ⊗ was the Kronecker product. The advantage of the multivariate approach was to explicitly model genetic covariances between time points.

Two methods for modeling environmental spatial variation were investigated, namely 2DSpl and AR1 models. The 2DSpl method created Z_p_ *u*_*p*_ as a function of the row (*r*) and column (*c*) position of the plot in the field and can be written as

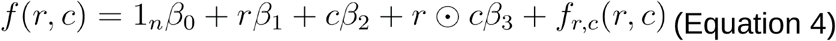

where ⊙ was the vector product and *f*_*r,c*_ (*r,c*) was a smoothing function (Rodríguez-Álvarez et al. 2018). The 2DSpl models were fitted using Sommer in R (R 3.6.3, Sommer 4.1.3) (Covarrubias-Pazaran 2016). In the univariate case, named 2DSplU, the variance of the spatial effect followed

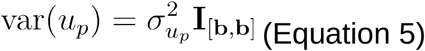

where I_[b,b]_ was an identity matrix with order equal to the total number of plots *b* in the experiment. In the multivariate case, named 2DSplM, the spatial variance was defined as

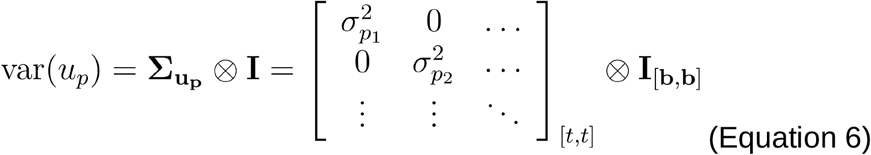

where the diagonal matrix 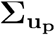 was of order [*t,t*] denoting the number of time points.

Contrastingly, the AR1 method explicitly defined a separable autoregressive covariance structure. In the univariate case, named AR1U, the spatial variance was defined as

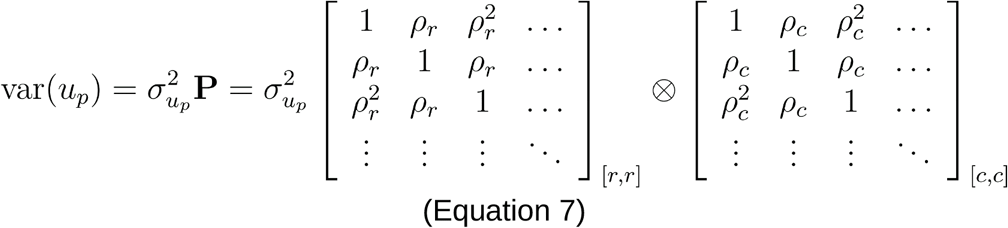

where P was a shorthand for the illustrated separable autoregressive structure, *ρ*_*r*_ and *ρ*_*c*_ were correlations among rows and columns, respectively, and [*r,r*,] and [*c,c*] indicated the dimensions for the matrices as the total number of rows and columns, respectively. With the AR1 model, experimental plots which were farther away had correlations which decreased exponentially on a unit-by-unit basis (e.g. *ρ* > *ρ* > ^2^ *ρ* >^3^…). In the multivariate case, named AR1M, the variance of the random spatial effect was written as

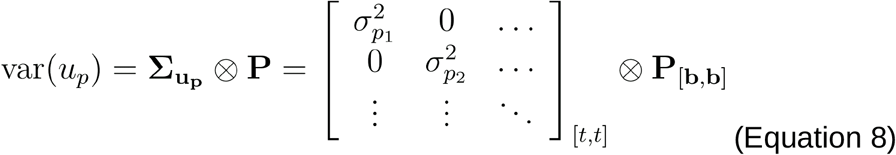

where the diagonal matrix 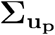 was of order [*t,t*] denoting the number of time points and the matrix *P* was the same as in the univariate case. The AR1 methods were fitted using ASReml-R (R 3.6.3, ASReml-R 4.1.0.126) (A. R. Gilmour et al. 2002). In the tested univariate and multivariate spatial models, the random residual error, *e*, followed 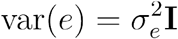 and 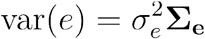, respectively, where 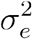 was the error variance and Σ_e_ was unstructured.

As an alternative to spatial mixed models, random regression (RR) models were explored for separating permanent environmental (PE) NLEE from additive genetic effects (Schaeffer 2004; Kirkpatrick, Lofsvold, and Bulmer 1990; Van der Werf, Goddard, and Meyer 1998; Arnold, Kruuk, and Nicotra 2019). PE offered a purely longitudinal representation of the spatial heterogeneity in the field. The RR model for a repeated measurement was written as

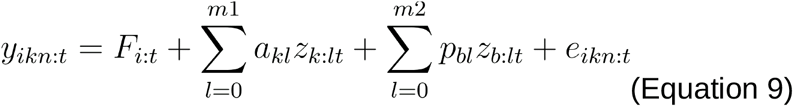

where *y*_*ikn:t*_ was the th observation on the *k*th genotype at time *t* belonging to the *i*th fixed factor. This study defined a single fixed effect, *F*_*i:t*_, for the replicate nested with the imaging event date. The variable *a*_*kl*_ represented the additive genetic effect for the *k*th genotype, while the variable *P*_*bl*_ denoted the PE effect for an experimental plot *b* planted in the field. The incidence covariables *z*_*k:lt*_ and *z*_*b:lt*_ mapped the random genetic and PE effects, respectively, to the observations. The variables *m*1 and *m*2 represented the total order of the random regression functions, and *e*_*ikn:t*_ was the heterogeneous random residual error.

The RR model was written in mixed model matrix notation as in Equation 1, however, the incidence matrices Z_a_ and Z_p_ are structured to account for the random regression covariance function coefficients, *u*_*a*_ and *u*_*p*_ and denoted the random additive genetic and PE regression coefficients, respectively. The overall variance was written as 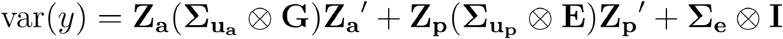; in this equation, G represented the GRM and E represented a plot-to-plot covariance matrix capturing environmental effects, ideally computed from envirotyping information. Envirotyping aimed to uniquely define the complete environment of an organism by including soil, climate, and developmental parameters; however, this study focused on applying NLEE (Xu 2016).

Notably different from the spatial mixed models described previously, the incidence matrices Z_a_ and Z_p_ can be structured to contain continuous variables representing time in the form of GDD for models using Legendre polynomials or linear spline functions. Also, the additive genetic variance for any hybrid at a specific time point 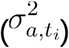 and the additive genetic covariance for any hybrid between two time points (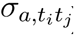) can be calculated using 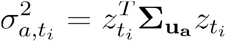 and 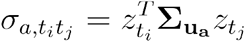, respectively, where 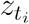 and 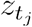 are vectors of the continuous random regression function evaluated at time points *t*_*i*_ and *t*_*j*_, respectively. Similar expressions for the PE variance 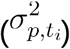 and covariance 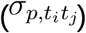 can be written as 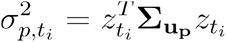 and 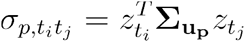, respectively. In this study, solutions to the RR model were found using the BLUPF90 family of programs (Misztal et al. 2002).

The RR model had computational benefits over the spatial mixed models described previously. Firstly, the number of variance components to estimate was equal 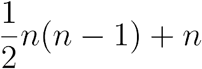 to, following the order *m*1 and *m*2 of the random regression functions, regardless of the number of time points *t* represented in the observations *y*. Secondly, continuous curves for the random additive genetic and PE effects could be evaluated for any time point because the random regression coefficients fit a covariance function. Thirdly, there was flexibility in the type of regression function that can be fitted, for instance, splines versus Legendre polynomials; in this study, third order Legendre polynomials were considered (Szeg 1939). Fourthly, flexible specification of E allowed envirotyping information to be accounted for in the model.

This study explored six different structures for the PE covariance matrix E. The first approach, named RRID, defined, E = I where I was the identity matrix. This approach treated all experimental plots as independent. The second approach, named RREuc, computed E using the inverse Euclidean distances between experimental plots and standardized between 0 and 1. This was written as 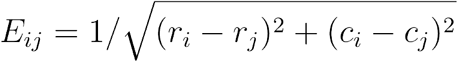, where *E*_*i,j*_ was an element of E denoting the relationship between plot and, and the variables *r*_*i*_, *c*_*i*_, *r*_*j*_, and *c*_*j*_ were the row and column positions of the plot, respectively. The inverse was used to give plots which were farther away from each other a smaller value than plots which were nearby. The third approach, named RRSoilEC, computed an empirical correlation matrix E from plot-level values of soil EC, dEC, and d2EC. The fourth approach, named RRSoilAlt, computed an empirical correlation matrix E from plot-level values of soil Alt, dAlt, and d2Alt. The fifth approach, named RR2DSpl, created an empirical correlation matrix E from the NLEE random spatial effects resulting from the 2DSplM model. The sixth approach, named RRAR1, created an empirical correlation matrix E from the NLEE random spatial effects resulting from the AR1M model.

The presented models allowed estimation of additive genetic and local environmental random effects; however, the true genetic and environmental effects were unknown to us. Therefore, to evaluate the robustness of the tested models and their ability to detect field environment features, six different simulation scenarios were conducted. The simulation methods are described in File S1. The simulations were designed to represent purely environmental effects in the field, such as due to soil heterogeneity, altitude, and soil elevation gradients. Six simulations named the Linear, 1D-N, 2D-N, AR1xAR1, and RD simulation, were each tested by varying the correlation across time to be 0.75, 0.90, and 1.00 and by setting the simulated variance to be 10%, 20%, and 30% of the total phenotypic variance.

Evaluating the accuracy of the simulation process was a five-step procedure. Firstly, for a target model, meaning one of the first-stage univariate/multivariate spatial mixed models or RR models, NLEE were separated from additive genetic effects present in the real NDVI HTP for a given field experiment. Secondly, the computed NLEE were subtracted from the NDVI HTP in order to minimize latent spatio-temporal effects in the NDVI. Thirdly, the target simulation values, meaning one of the six simulation processes, were scaled between 0 and 1, then subsequently scaled to account for either 10%, 20%, or 30% of the observed NDVI phenotypic variation, and finally were added onto the minimized NDVI HTP. Fourthly, the target model computed LEE for the simulation-adjusted NDVI HTP. Fifthly, the recovered LEE were correlated against the true target simulation values, returning prediction accuracy.

### Agronomic Genomic Prediction

The second question in this study was whether the longitudinal NDVI or NLEE data could be used to improve spatial corrections, specifically for genomic prediction (GP) of agronomic traits. Firstly, the following baseline models were defined under the GBLUP framework. The baseline GBLUP model was written as

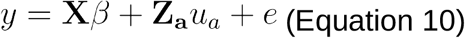

where *y* was the agronomic trait of interest, *β* was for fixed effects like replication,*u*_*a*_ was the random additive genetic effect of the hybrid, X and Z_a_ were incidence matrices linking effects to *y*, and *e* was the residual error. The variance of the random additive genetic effect was defined as 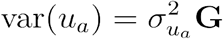 where G was the GRM and 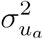 was the additive genetic variance. The error variance was defined as 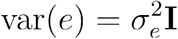 where 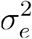 was the residual variance. Control GBLUP models named G, G+2DSpl, and G+AR1, written respectively as

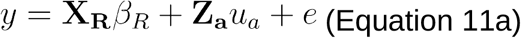

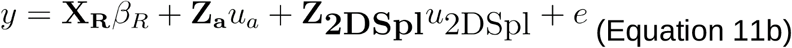

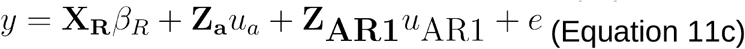

were used as baseline models for agronomic genomic prediction and *β*_*R*_ was a fixed effect for replication linked to corresponding incidence matrix X_R_. Equation 11a was the simplest GBLUP case. Incorporating spatial corrections, Equations 11b and 11c added 2DSpl and AR1 random effects, respectively, to account for the row and column positions of the plots in the field on which the agronomic trait was measured. Equation 11b followed the 2DSpl definition from Equation 4, while Equation 11c followed the AR1 definition from Equation 7. Importantly, these three baseline models did not leverage information from the first stage or from the aerial image HTP measurements.

A final baseline defined the multi-trait model (M) in Equation 12,

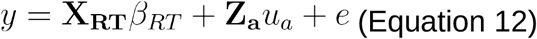

where *y* was a vector of both the HTP NDVI time points and the agronomic trait (e.g. GY or EH or GM), *β*_*RT*_ was a fixed effect for replicate nested with trait, *u*_*a*_ was a vector of the random additive genetic effects for all traits, and *e* was the random residual variance. The random additive genetic (*u*_*a*_) and residual (*e*) covariances were unstructured across the traits, as in Equation 3. Given the difficulty of fitting large numbers of traits in M, only the two HTP NDVI timepoints with the highest correlation to GY were included in *y*.

Secondly, to improve on the baseline GBLUP models first-stage NLEE were integrated into the second stage following two implementations. The first implementation modeled the NLEE as a plot-to-plot covariance structure for random effects (L), following:

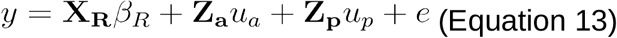

The variance of the random plot effects *u*_*p*_ followed 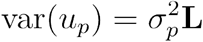, where L was an empirical matrix derived from correlating the NLEE for all plots across time points.. This model was similar to Equation 11c in which the AR1 process explicitly defined a plot-to-plot covariance structure; however, rather than define distance-based assumptions, Equation 13 utilized observed spatial heterogeneity. The second column of Table 2 summarized the tested L two-stage model names in relation to the first-stage.

**Table 2:**
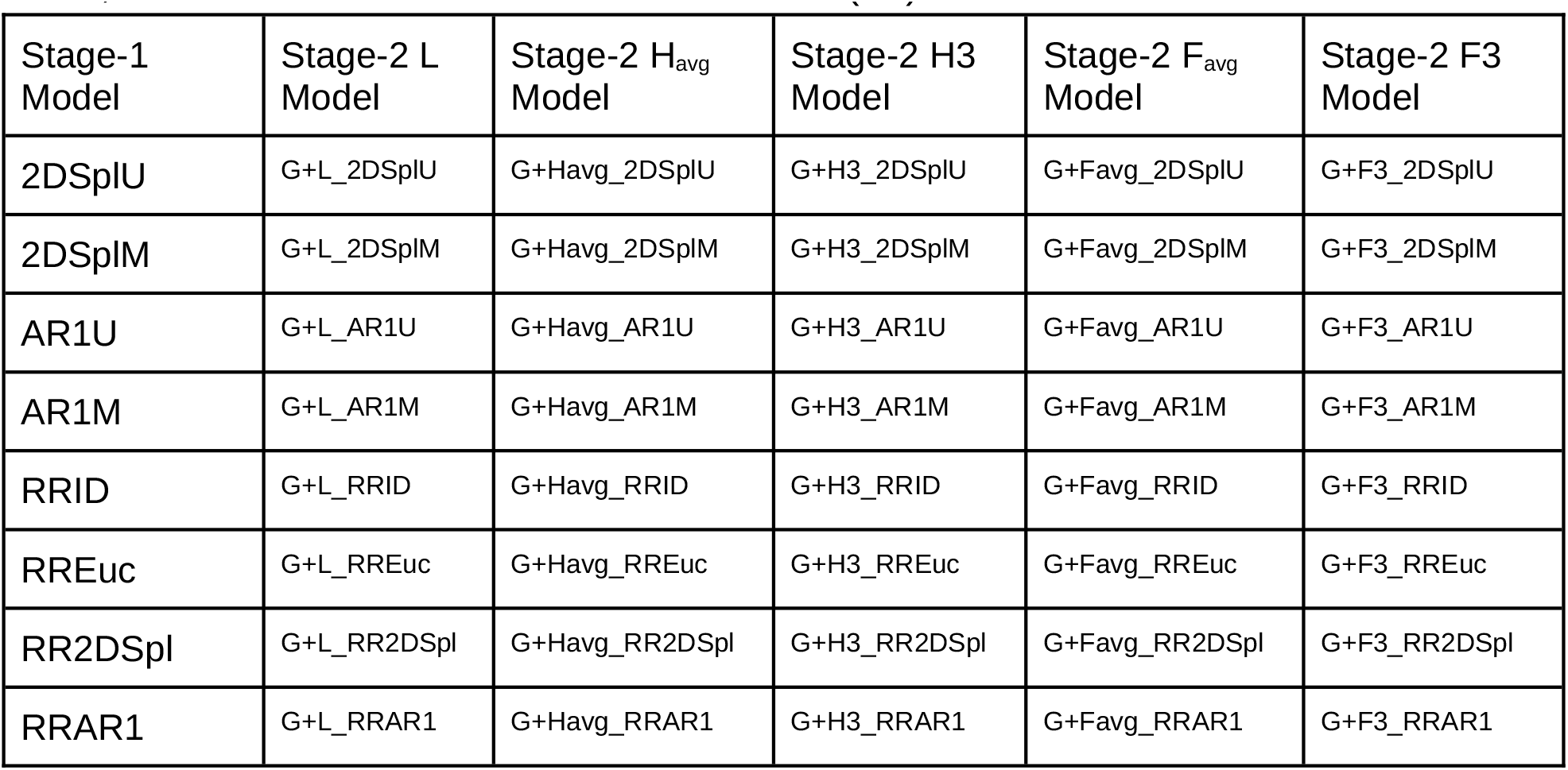
Listed are all non-soil two-stage models tested in this study. The first column listed the first-stage models used to separate additive genetic effects from NDVI local environment effects (NLEE). Subsequent columns listed models where the second-stage was implemented as a plot-to-plot covariance (L), as a continuous average fixed effect (Havg), as three distinct continuous fixed effects (H3), as a binned average fixed effect, and as three distinct binned fixed effects (F3).

Alternatively, fixed factor effects (FE) were defined and followed four variations:

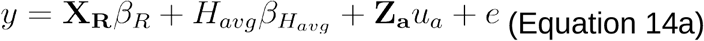

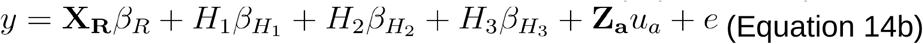

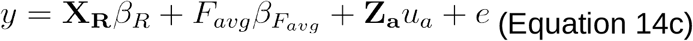

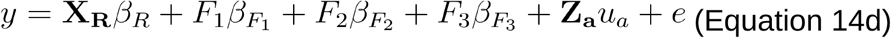

In Equation 14a the fixed factor *H*_*avg*_ performed a continuous regression on the average first-stage NLEE across all time points, accounting for fixed effects 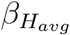. Alternatively, Equation 14b performed a continuous regression on the fixed factors *H*_1_, *H*_2_, and *H*_3_ by splitting time points into the previously defined P0, P1, and P2 growth phases, respectively, and then averaging the first-stage NLEE within each. This model accounted for fixed effects 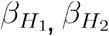 and 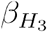 for the P0, P1, and P2 growth phases, respectively. Equation 14c and 15d used binned fixed effects derived by reassigning the first-stage NLEE to quartile factors (1=0-25%, 2=26-50%, 3=51-75%, 4=76-100%), representing poor, marginal, good, and high performing levels. The *F* _*avg*_ fixed factor in Equation 14c was computed by averaging over all time points, while the *F*_1_, *F*_2_, and *F*_3_ factors in Equation 14d were computed by splitting the time points into the P0, P1, and P2 growth phases, respectively, and then averaging within each. Columns 3 to 6 of Table 2 summarize the tested FE two-stage model names in relation to the first-stage.

Soil altitude and EC were measured in 2019, enabling Equations 13, 14a, and 14c to account for soil information rather than first-stage NLEE. It was not possible to test Equation 14b or 14d with soil information because only a single soil measurement was collected. Representing Equation 13, a model named G+L_RRSoilEC constructed *L* by correlating soil EC, dEC, and d2EC, while a model named G+L_RRSoilAlt constructed *L* by correlating soil Alt, dAlt, and d2Alt. Using soil measurements in Equation 14a, models named G+Havg_Soil_Alt, G+Havg_Soil_dAlt, G+Havg_Soil_d2Alt, G+Havg_Soil_EC, G+Havg_Soil_dEC, and G+Havg_Soil_d2EC represented *H*_*avg*_ derived from the plot-level soil Alt, dAlt, d2Alt, EC, dEC, and d2EC, respectively. Whereas for Equation 14c, models named G+Favg_Soil_Alt, G+Favg_Soil_dAlt, G+Favg_Soil_d2Alt, G+Favg_Soil_EC, G+Favg_Soil_dEC, and G+Favg_Soil_d2EC represented *F*_*avg*_ derived from the plot-level soil Alt, dAlt, d2Alt, EC, dEC, and d2EC, respectively.

Model performance was tested by measuring heritability, model fit, and genotypic effect estimation across replicates. Narrow-sense genomic heritability *h*^2^ was defined as 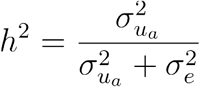 where 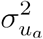 was the additive genetic variance and 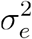 was the residual error variance. Heritability gives insight into the degree to which additive genetic effects are driving phenotypic variation over residual error. Model fit was defined as the correlation between fitted model predictions *ŷ* and the true phenotypic values *y*, written as cor(*ŷ,y*). Finally, genotypic effect estimation across replicates, written as cor(*g*_Rep1_ *g*_Rep2_), involved first partitioning the agronomic trait datasets by replicate, then running the target model on each replicate, and finally correlating the random genetic effect estimates,, across the replicates. The field experiments in this study lent themselves to testing cor(*g*_rep1_,*g*_Rep2_), because two contiguous replicates were designated in all fields. Genotypic effect estimation across replicates was the primary model metric in this study because it is an important indicator for whether a spatial correction has effectively accounted for field heterogeneity.

## Results and Discussion

### Local Environmental Effects

Before turning attention to the NDVI HTP, spatial corrections for the agronomic traits of grain yield (GY), grain moisture (GM), and ear height (EH) were computed. As defined in Equation 11b, Figure 1 illustrates heatmaps of the 2DSpl spatial effects over the rows and columns of the experimental plots in the 2017_NYH2, 2019_NYH2, and 2020_NYH2 field experiments. Each year the trial was planted in a distinct field location. The heatmaps resolved major, poorly performing regions for GY and EH centered around the row-column positions of (35,2), (19,8), and (25,6) in 2017_NYH2, 2019_NYH2, and 2020_NYH2, respectively. For all three experiments, an inverse spatial pattern was visible for GY and GM, while a similar spatial pattern was visible for GY and EH. The same pattern between traits was evidenced in 2015_NYH2, illustrated in Figure S7. In 2015_NYH2 a poor performing region for GY and EH was centered near the row-column position of (88,9). In all four years, the proportion of phenotypic variation explained by the 2DSpl spatial effect ranged from +/-25 bu/acre of GY, +/-2% of GM, and +/-15 cm of EH. These results illustrated the importance of spatial heterogeneity on the agronomic traits.

**Figure 1:**
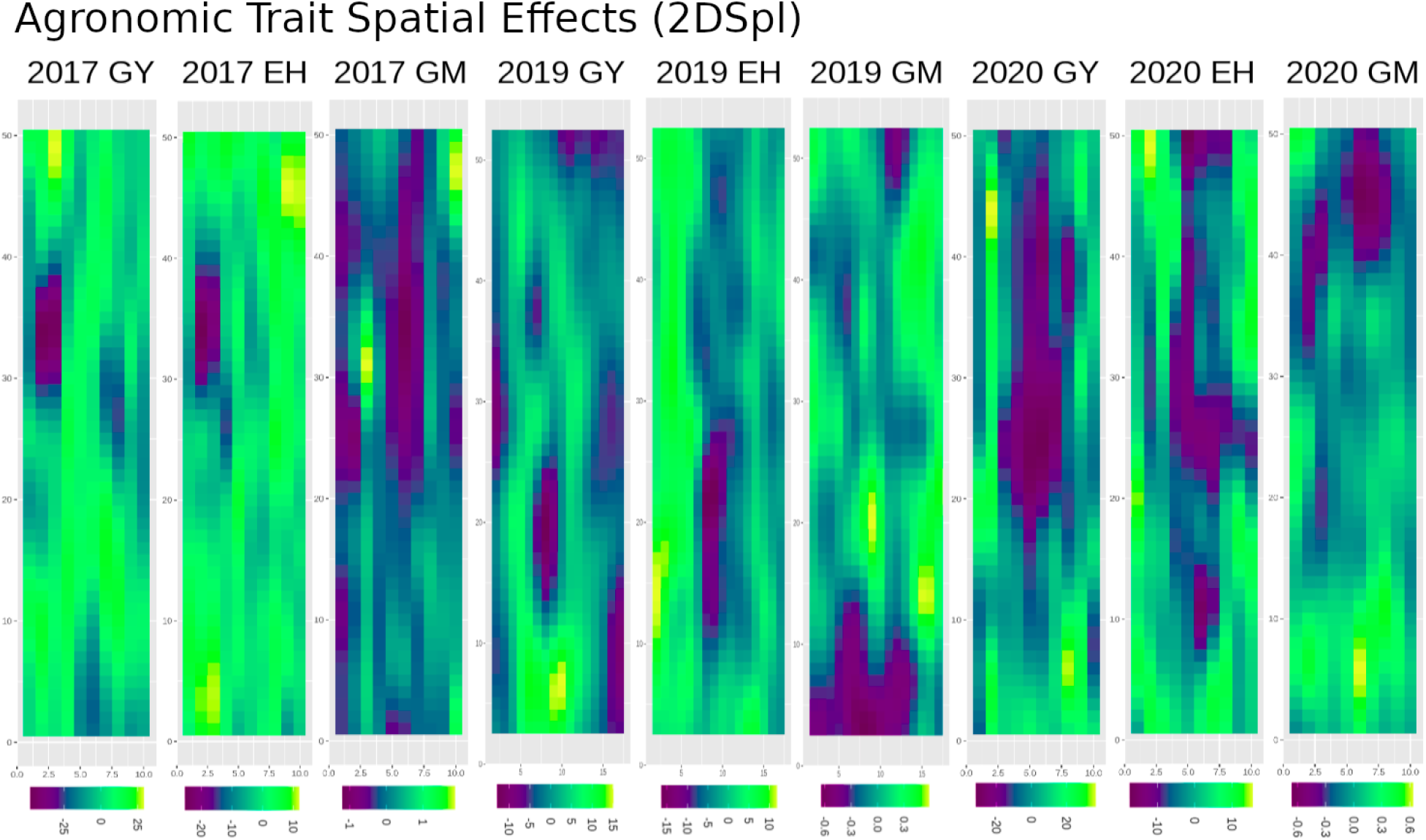
Two-dimensional spline (2DSpl) spatial random effects detected in the agronomic traits of grain yield (GY), grain moisture (GM), and ear height (EH) in the 2017, 2019, and 2020 field experiments. The 2015 experiment is shown in Figure S7.

Similar patterns of spatial random effects were found using the 2DSpl and AR1 models defined in Equations 11b and 11c, respectively. Table 3 lists the correlations between the 2DSpl and AR1 spatial effects for GY, GM, and EH in the 2017_NYH2, 2019_NYH2, and 2020_NYH2 experiments. The strongest average correlation was 0.88 for EH, followed by 0.87 for GY, and 0.49 for GM. The 2015_NYH2 experiment was not included in Table 3 because the AR1 model did not converge.

**Table 3:**
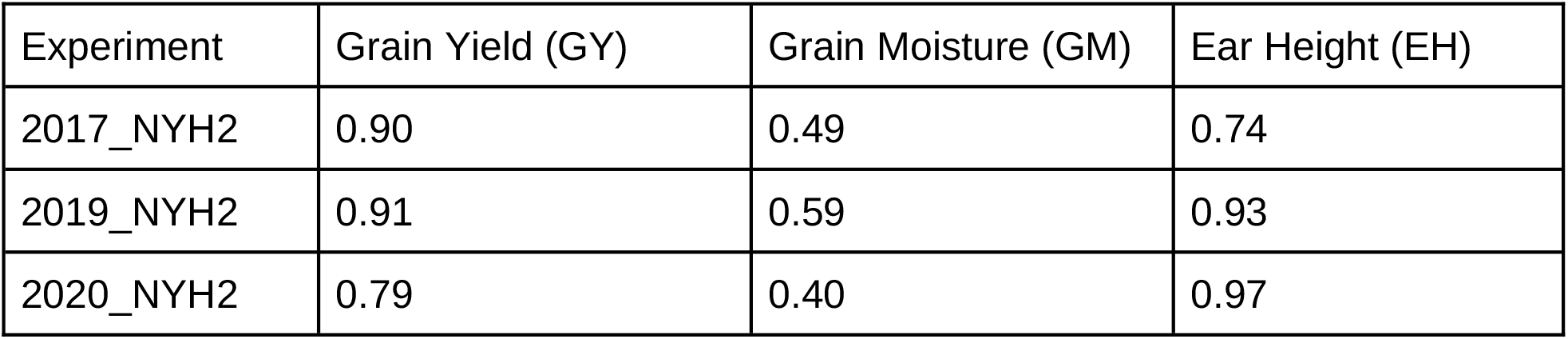
Correlations between 2DSpl and AR1 spatial effects for GY, GM, and EH in the 2017, 2019, and 2020 field experiments.

To understand correspondence between LEE affecting NDVI and the end-of-season agronomic traits, first-stage NLEE were compared to GY, GM, and EH spatial effects. Figure 2 shows correlations between the 2DSpl GY spatial effects and the 2DSplU NLEE across 12 time points in the 2020_NYH2 field experiment. Illustrated in Figure 2, correlations at 38 and 48 days after planting (DAP) were 0.6 and 0.7, respectively, and correlations fluctuated between 0.5 and 0.7 throughout the season. Corresponding heatmaps in Figure 2 illustrate spatial distributions over the rows and columns of the experimental plots, and consistently reveal a large region near the center of the field negatively impacting both GY and NDVI. For reference, phenotypic correlations between NDVI and GY in 2020_NYH2 ranged from a low of 0.04 at 133 DAP to a high of 0.39 at 54 DAP, which were weaker than the correlations observed in Figure 2. Similarly, the random regression (RR) model PE effects correlated with the GY spatial effects more strongly than the NDVI and GY themselves. Figure S8 illustrates RRID PE effects correlated against GY and the GY 2DSpl and AR1 spatial effects. The RRID PE tended to be strongly correlated through time and identified similar spatial patterns as the spatial effect models.

**Figure 2:**
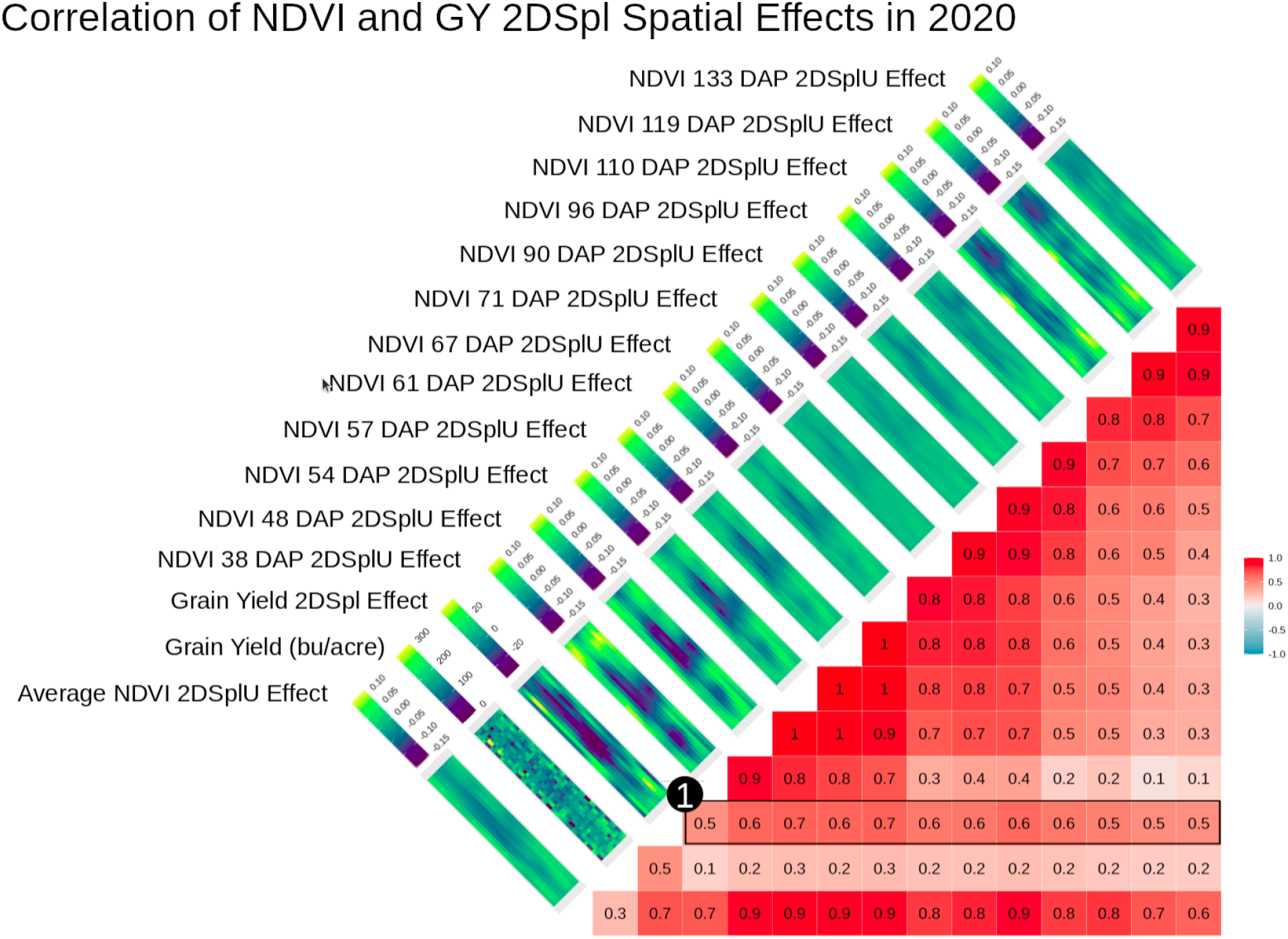
2020_NYH2 2DSplU NLEE observed over 12 time points correlated with GY and GY 2DSpl spatial effects. Corresponding heatmaps showed values over the rows and columns of all experimental plots in the field and revealed similar spatial patterns. As indicated by (1), correlations of 0.5 to 0.7 between GY 2DSpl and the 2DSplU NLEE were found throughout the growing season, even early on at 38 DAP.

In 2019_NYH2, 2015_NYH2, and 2017_NYH2 similar correlations between 2DSplU and GY spatial effects were observed, as illustrated in Figure 3, Figure S9, and Figure S10, respectively. For reference, phenotypic correlations between NDVI and GY in 2015_NYH2, 2017_NYH2 and 2019_NYH2 ranged from a low of 0.13 at 126 DAP, 0.17 at 25 DAP, and 0.33 at 110 DAP, respectively, to a high of 0.42 at 92 DAP, 0.49 at 91 DAP, and 0.67 at 84 DAP, respectively. Therefore, in all experiments the NLEE were more correlated to the GY 2DSpl effects across the growing season than the NDVI were correlated to GY. Furthermore, in all years, the spatial patterns affecting GY and EH were detectable by NLEE to a large degree (> 0.5 correlation), evidenced early on in the growing season at 76 DAP or less.

**Figure 3:**
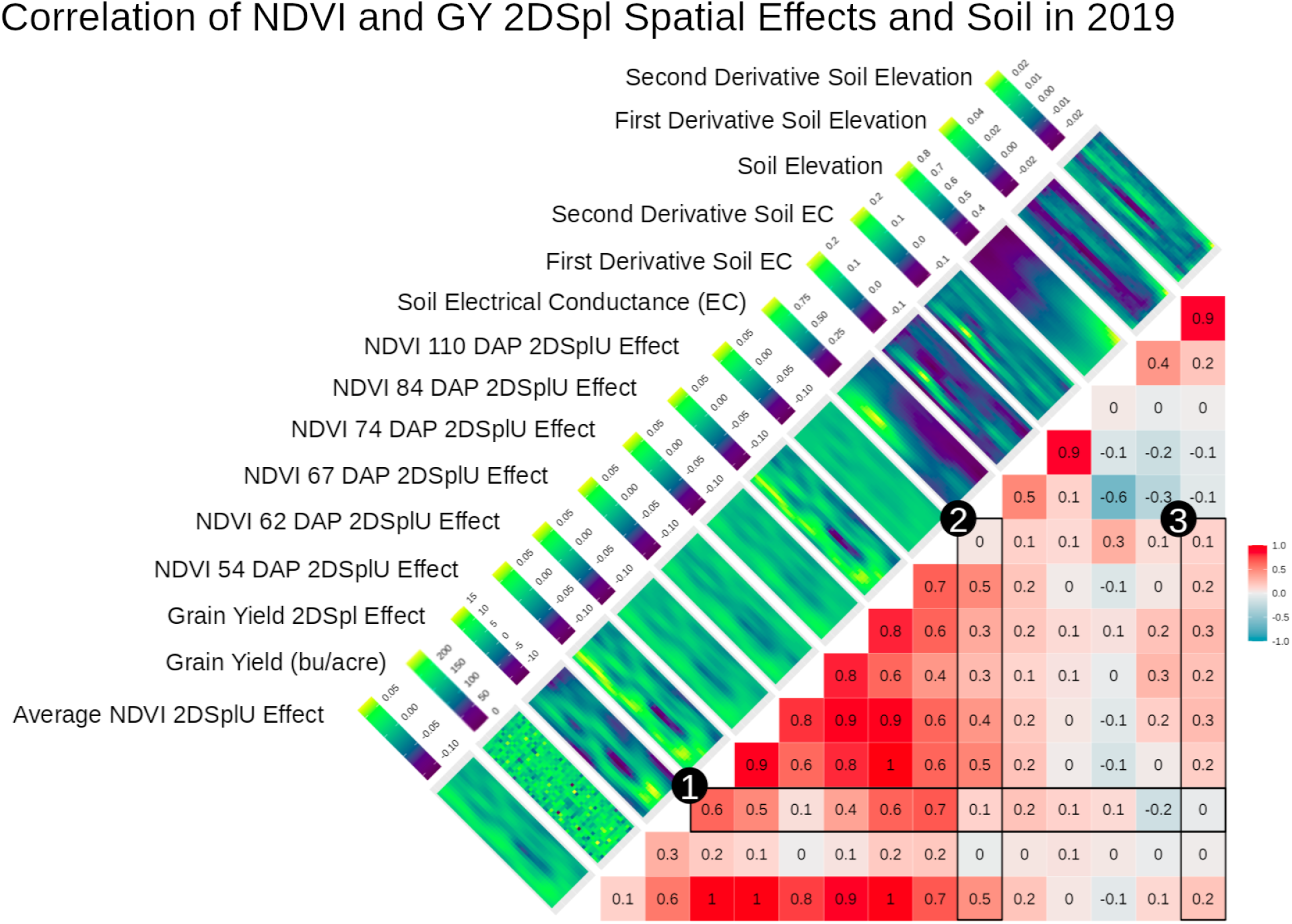
2019_NYH2 2DSplU NLEE observed over 6 time points correlated with GY and GY 2DSpl spatial effects. Corresponding heatmaps showed values over the rows and columns of all experimental plots in the field and revealed similar spatial patterns. As indicated by (1), correlations of 0.1 to 0.7 between GY 2DSpl and the 2DSplU NLEE were found throughout the growing season. The strongest correlation of 0.7 was observed at 110 DAP, however, at 54 DAP a correlation of 0.6 was observed. Soil EC and Alt, as well as the first and second numerical two-dimensional derivatives, were included against the NLEE and the following were observed: (2) correlations up to 0.5 for EC and (3) correlations up to 0.3 for d2Alt. The AR1 analog in Figure S11 showed that the AR1U NLEE were less correlated with the soil parameters than the 2DSplU NLEE, and the AR1U NLEE were less correlated with GY AR1 effects than the 2DSplU NLEE were correlated with the GY 2DSpl effects.

As a potential alternative to aerial imaging and to better understand the observed spatial effects, soil EC, Alt, and the first and second two-dimensional numerical derivatives (dEC, d2EC, dAlt, d2Alt) were compared with the NLEE in the 2019_NYH2 field experiment. Figure 3 includes correlations and heatmaps of the soil measurements with the spatial effects and demonstrated: (1) the 2DSpl spatial effects of GY in 2019_NYH2 correlated up to 0.7 with 2DSplU NLEE, (2) soil EC correlated up to 0.5 with 2DSplU NLEE, and (3) soil d2Alt correlated up to 0.3 with 2DSplU NLEE. In Figure 3, the soil elevation gradients, represented by heatmaps of Alt, EC, and their derivatives, highlighted the contours of the observed 2DSpl spatial effects. Figure S11 presents an analog to Figure 3 showing AR1U NLEE, GY, and GY AR1 spatial effects in 2019_NYH2, and showed (1) overall weaker correlations between the GY AR1 spatial effects and the AR1U NLEE, with a high of 0.5, (2) similar correlations to soil EC with a high of 0.5, and (3) overall weaker correlations to soil d2Alt with a high of 0.2. Therefore, the AR1U NLEE were less correlated with the soil parameters than the 2DSplU NLEE, and the AR1U NLEE were less correlated with GY AR1 effects than the 2DSplU NLEE were correlated with the GY 2DSpl effects.

Table 4 summarizes correlations between the soil information in 2019_NYH2 and the model NLEE, illustrating how the average first-stage NLEE across all time points correlated to soil EC, dEC, d2EC, Alt, dAlt, and d2Alt. Table 4 indicates 2DSplU had the strongest correlation to EC of 0.46 and also correlated relatively strongly with d2Alt. The d2Alt tended to correlate more strongly than Alt or dAlt with the NLEE, indicating the importance of elevation gradients in the field. The correlations between 2DSpl NLEE and soil parameters indicated that soil information was capturing similar spatial information as the NDVI aerial imaging.

**Table 4:** Correlations between average NLEE in 2019_NYH2 computed using different first-stage models and the soil EC, dEC, d2EC, Alt, dAlt, and d2Alt measurements.

| Model | Soil EC | Soil dEC | Soil d2EC | Soil Alt | Soil dAlt | Soil d2Alt |
| --- | --- | --- | --- | --- | --- | --- |
| 2DSplU | 0.46 | 0.21 | 0.04 | -0.06 | 0.11 | 0.21 |
| 2DSplM | 0.21 | 0.12 | 0.06 | 0.08 | 0.34 | 0.29 |
| AR1U | 0.39 | 0.18 | 0.04 | -0.06 | 0.06 | 0.16 |
| AR1M | 0.01 | 0.04 | 0.06 | 0.10 | 0.25 | 0.21 |
| RRID | 0.10 | 0.06 | 0.04 | 0.04 | 0.14 | 0.17 |
| RREuc | -0.06 | 0.03 | 0.09 | 0.07 | 0.14 | 0.15 |
| RRAR1 | -0.18 | -0.06 | 0.03 | 0.11 | 0.23 | 0.21 |
| RR2DSpl | -0.19 | -0.07 | 0.02 | 0.13 | 0.24 | 0.23 |
| RRSoilEC | -0.22 | -0.08 | 0.03 | 0.12 | 0.25 | 0.22 |
| RRSoilAlt | -0.21 | -0.09 | 0.04 | 0.12 | 0.25 | 0.23 |

Summarizing correlations between the agronomic trait 2DSpl effects and the tested first-stage model LEE, Figure 4 presents results for all agronomic traits and years. Figure 4 indicates the 2DSplU, 2DSplM, and AR1U models produced NLEE most correlated to the 2DSpl effects of GY and EH in all years and in nearly all time points, while the RR2DSpl, RRAR1, and RRSoilEC models were most correlated to the 2DSpl effects of GM in 2015, 2017, and 2019. The 2DSplU and AR1U NLEE correlated with GY and EH 2DSpl spatial effects greater than 0.5 in all years at 80 to 90 DAP. Significant similarities were seen between models run on traits within a given year, for instance both GY and EH in 2017 showed a large peak at 110 DAP and in 2020 both showed a continuous gradual decline. There was an inverted behavior between the GY and GM spatial effects in all years, describable as: in 2015 a high for GY and a low for GM at 105 DAP, in 2017 a high for GY and a low for GM at 110 DAP, in 2019 a high for GY and a low for GM around 90 to 100 DAP, and in 2020 a gradual decline for GY and a gradual incline for GM. In contrast, there was a similar behavior between the GY and EH spatial effects in all years. Figure S12 illustrates the AR1 analog of Figure 4 with agronomic trait AR1 spatial effects instead of 2DSpl effects and demonstrated weaker correlations to the NLEE in all traits and all years. Highly similar patterns in the correlation curves were observed, however, there was a tendency for the GM AR1 spatial effects to correlate with RRID, RR2DSpl, and RRAR1 NLEE more strongly than the 2DSpl or AR1 NLEE.

**Figure 4:**
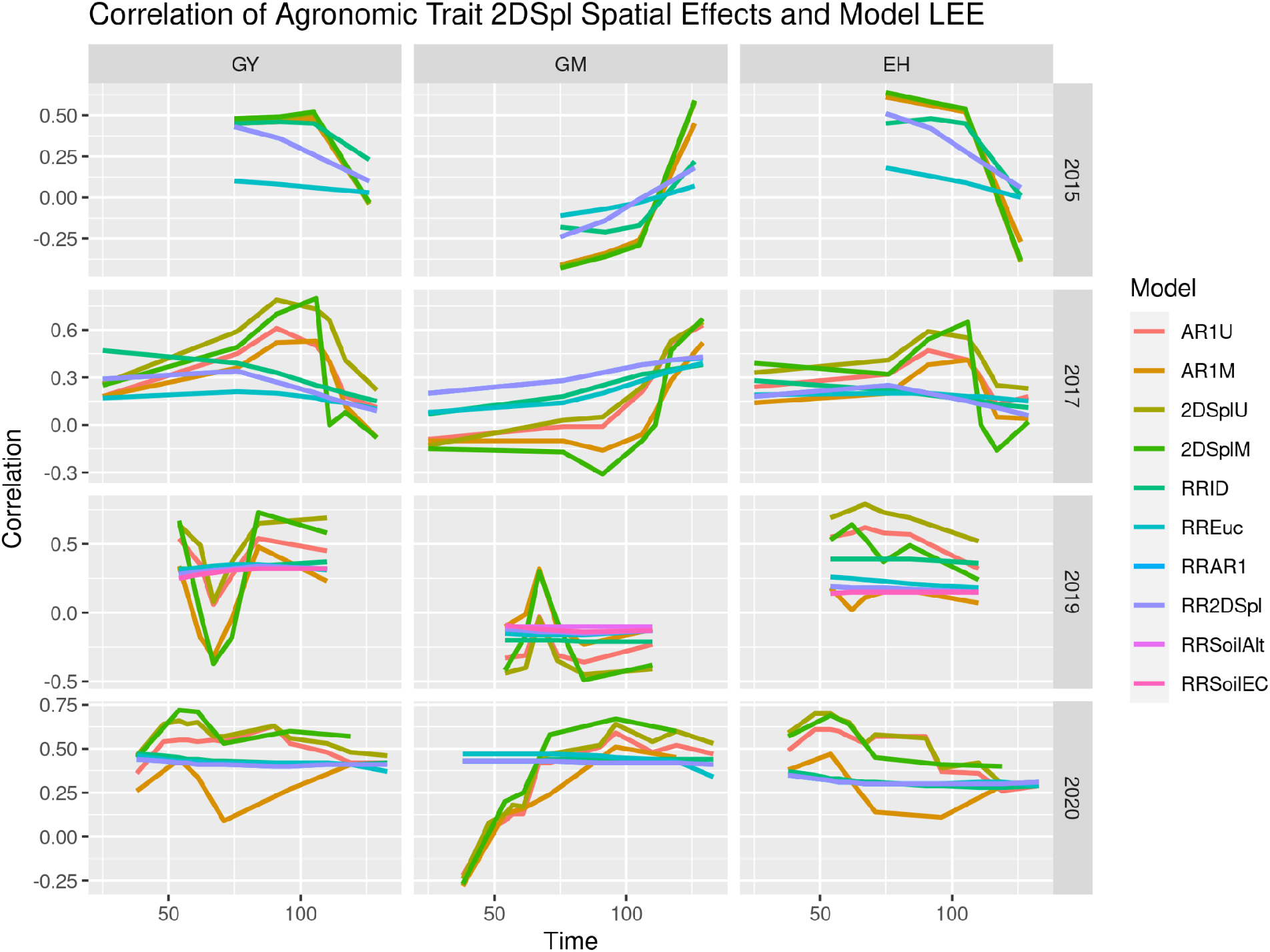
Correlations between 2DSpl spatial effects of agronomic traits and LEE from the tested first-stage models. Spatial effects for GY, GM, and EH in the 2015_NYH2, 2017_NYH2, 2019_NYH2, and 2020_NYH2 field experiments were compared with model LEE across the growing season. Models including soil information rather than NDVI in the first-stage, named RRSoilAlt and RRSoilEC, were also included for 2019_NYH2. The AR1 analog in Figure S12 illustrated similar patterns, but with weaker correlations.

Simulation tested the efficacy of the first stage in detecting known environmental field effects. In each of the six simulation processes (linear, 1D-N, 2D-N, AR1xAR1, random, and RD) ten iterations were performed, each time generating a new simulation. The simulated environmental variance was tested at 0.1, 0.2, and 0.3 times the proportion of phenotypic variation, and the correlation between time points was tested at 0.75, 0.90, and 1. Figure S22, Figure S23, and Figure S24 illustrate the results for all simulation scenarios using 2017_NYH2, 2019_NYH2, and 2020_NYH2 NDVI phenotypes, respectively, demonstrating the impacts of varying the simulation environmental variance as well as the correlation of simulated environmental effects across the growing season. Increasing the variance tended to slightly increase the prediction accuracy, while decreasing the correlation between time points tended to decrease prediction accuracy. Figure 5 aggregates the prediction accuracies for the linear, 1D-N, 2D-N, AR1xAR1, and RD simulation scenarios and illustrated model groupings determined by a Tukey Honest Significant Difference (HSD) test.

**Figure 5:**
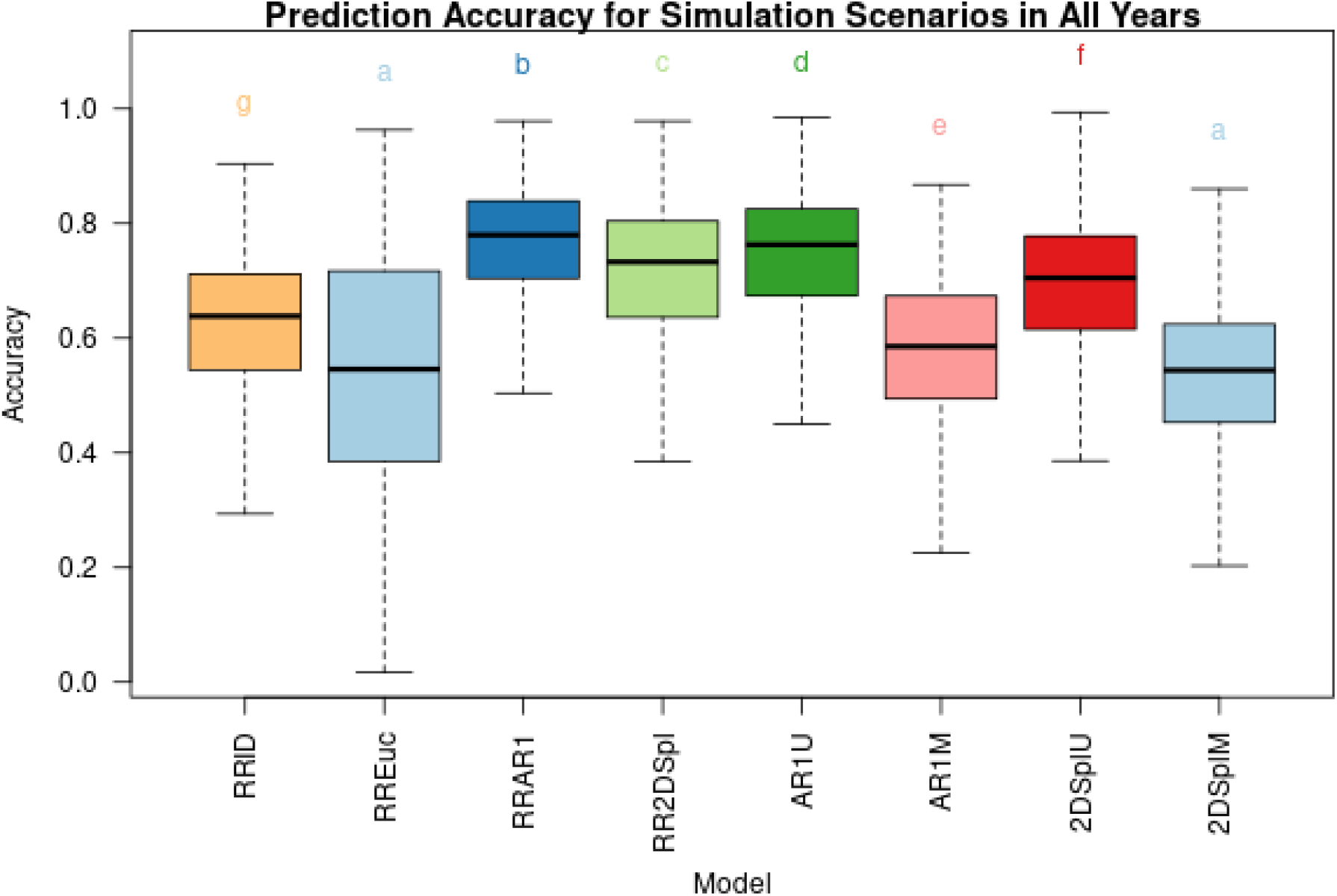
Aggregated prediction accuracy of the tested first-stage models (RRID, RREuc, RRAR1, RR2DSpl, AR1U, AR1M, 2DSplU, and 2DSplM) for the linear, 1D-N, 2D-N, AR1xAR1, and RD simulation scenarios using the 2017_NYH2, 2019_NYH2, and 2020_NYH2 NDVI data. Prediction accuracy is the correlation of the simulated environmental effect and the model’s recovered environmental effect. Models are grouped together after performing a Tukey HSD test.

The RREuc model showed relatively poor performance, possibly due to a mismatch in the geometry of the experiment because in reality the plots in the field were rectangular (e.g. 10ft by 3ft) and not perfectly square. The RREuc model was also most sensitive to the tested years and to changes in simulated variance and correlation. The AR1U model performed best in the AR1xAR1 scenario, while the 2DSplU model performed well in the Linear scenario. Specifying the PE covariance matrix allowed the RRAR1 and RR2DSpl models to perform consistently well in the 1D-N and 2D-N scenarios; however, by making no assumptions to the spatial structure the RRID model performs on average less than 10% worse and with comparable consistently.

### Agronomic Genomic Prediction

First-stage NLEE were incorporated into the second stage of the proposed GP approach using two distinct implementations, either modeling L or FE. Figure S13 illustrates genomic heritability, model fit, and genotypic effect estimation across replicates (cor(*g*_rep1_, *g*_Rep2_)) in the four years for GY, GM, and EH for all two-stage models when modeling L random effects. Baseline and spatially corrected GBLUP models, named G, G+2DSpl, and G+AR1 representing Equations 11a, 11b, and 11c, respectively, are shown. Statistical significance of the spatial corrections and two-stage models was compared to the baseline G model using a paired t-test. The best models were determined by ranking the t-test p-value of GY cor(*g*_Rep1_,*g*_Rep2_). The best five two-stage L models, representing Equation 13, were named G+L_AR1U, G+L_AR1M, G+L_2DSplU, G+L_2DSplM, and G+L_RRID.

Alternatively, modeling NLEE as FE followed four distinct definitions in Equations 14a, 14b, 14c, and 14d. Figure S17 illustrates genomic heritability, model fit, and cor(*g*_rep1_, *g*_Rep2_)in the four years for GY, GM, and EH for all two-stage models when modeling FE. The baseline and spatially corrected GBLUP models, named G, G+2DSpl, and G+AR1, respectively, are shown. As before, the best two-stage FE models were determined by ranking the t-test p-value of GY cor(*g*_Rep1_,*g*_Rep2_). The top eight models were named G+Havg_AR1U, G+Havg_2DSplU, G+Havg_2DSplM, G+Favg_2DSplU, G+H3_2DSplM, G+H3_RRID, G+F3_AR1U, and G+F3_2DSplU.

To observe performance of the proposed two-stage approach over the baseline G model, a difference (G Diff) was computed within each of the four years for heritability, model fit, and cor(*g*_Rep1_,*g*_Rep2_). Figure 6 illustrates G Diff for the baseline G+2DSpl and G+AR1 spatial correction models and for the best six models defining L (G+L) and FE (G+H). Figure S14 and Figure S18 illustrate all two-stage models when defining L and FE, respectively. The spatially corrected baseline models, G+2DSpl and G+AR1, demonstrate improvements over G. The G+2DSpl model provided significant improvements in heritability and model fit for all traits, and a significant improvement in GY cor(*g*_rep1_, *g*_Rep2_), while, G+AR1 provided significant improvements in model fit for all traits, and a significant improvement in GY heritability and cor(*g*_Rep1_,*g*_Rep2_). Increased GY heritability and cor(*g*_rep1_, *g*_Rep2_) demonstrated the value of performing spatial corrections; however, increased model fit may indicate overfitting.

**Figure 6:**
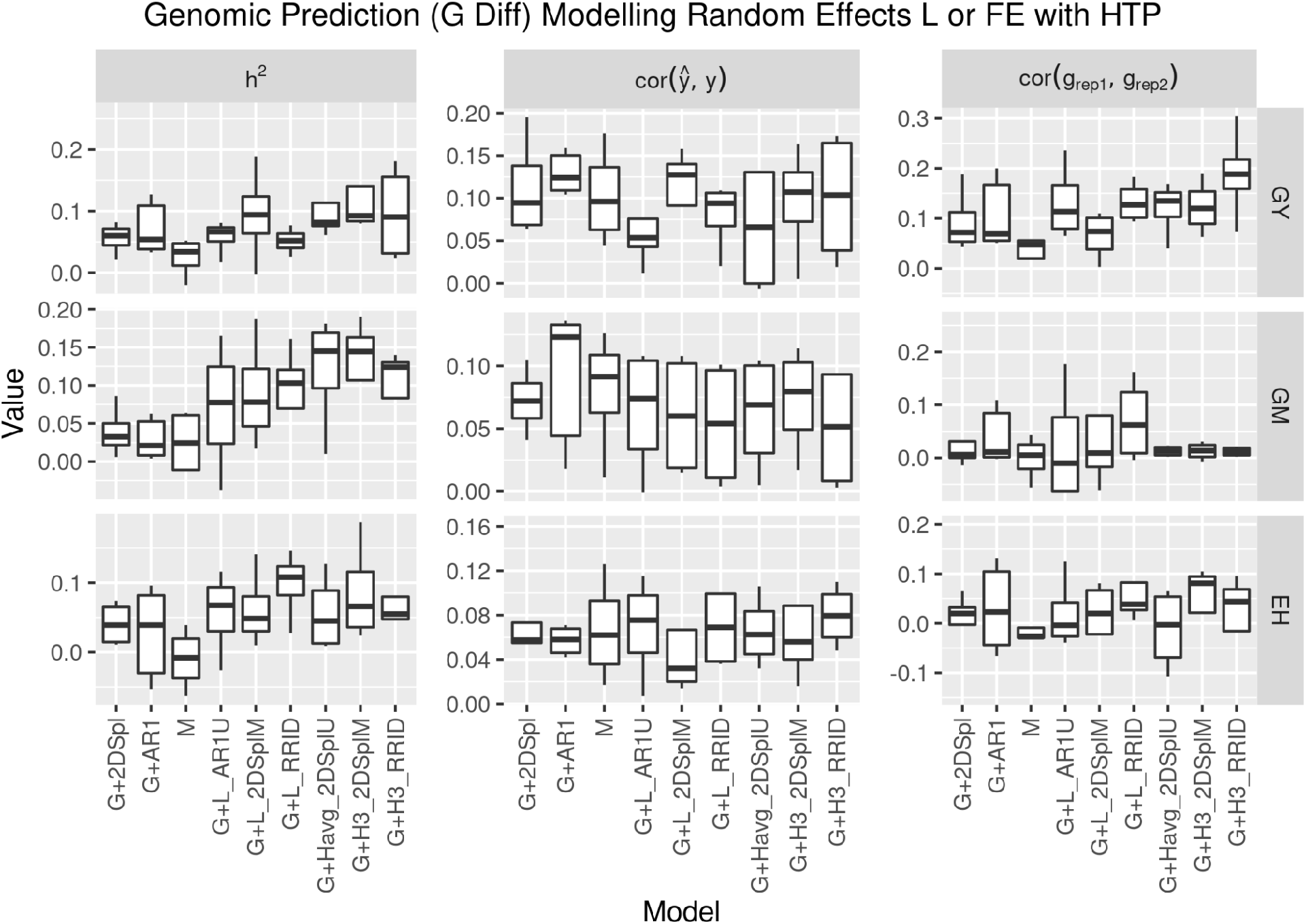
Differences compared to G (G Diff) for genomic heritability (*h*^2^), model fit (cor(*ŷ,y*)), and genotypic effect estimation across replicates (cor(*g*_rep1_, *g*_Rep2_))for GY, GM, and EH in the four years. The models G, G+2DSpl, and G+AR1 were baseline GBLUP and spatially corrected models, respectively, and M was a baseline multi-trait model. Illustrated are the best three two-stage models using L (G+L) and the best three two-stage models using FE (G+H), determined by ranking cor(*g*_rep1_, *g*_Rep2_) t-test p-value compared to G. The G+L and G+H models have L and FE, respectively, defined using NLEE of corresponding names.

Figure 6 demonstrates further improvements for the two-stage models over the baseline G, G+2DSpl, and G+AR1 models. Two-stage models incorporating NLEE improved heritability and cor(*g*_rep1_, *g*_Rep2_) for GY, GM, and EH more than the baseline spatially corrected models, and in addition, G+2DSpl and G+AR1 tended to increase model fit equally or greater than the two-stage models. The best two-stage models translated increased heritability to an increase in cor(*g*_rep1_, *g*_Rep2_), and avoided an inflation in heritability due to decreased residual error and increased model fit. Improvements to GY cor(*g*_rep1_, *g*_Rep2_) over baseline G, G+2DSpl, and G+AR1 models were summarized in Table 5 for the best six two-stage models when incorporating NLEE. The Table 5 columns of “Δ G”, “Δ G+2DSpl”, and “Δ G+AR1’’ indicate the mean and standard deviations of model differences in GY cor(*g*_rep1_, *g*_Rep2_) for the four years compared to G (G Diff), G+2DSpl (G+2DSpl Diff), and G+AR1 (G+AR1 Diff), respectively. Figure S15 and Figure S19 illustrate the G+2DSpl Diff for all models when defining L and FE, respectively. Figure S16 and Figure S20 illustrate the G+AR1 Diff for all models when defining L and FE, respectively. While GY and EH cor(*g*_rep1_, *g*_Rep2_) was improved, none of the FE models significantly improved GM cor(*g*_rep1_, *g*_Rep2_), a result potentially attributable to the lower correlations between NLEE and GM spatial effects seen in Figure 4 and Figure S12, the difficulty in detecting GM spatial effects seen in Table 3, and the small GM spatial variation seen in Figure 1.

**Table 5:** The best two-stage models versus the baseline GBLUP models (G, G+2DSpl, G+AR1) when comparing the correlation of genotypic effects across replicates cor(*g*_rep1_, *g*_Rep2_) for grain yield (GY). The best six two-stage models were listed, three defining L (G+L) and three defining FE (G+H). The columns “Δ G”, “Δ G+2DSpl”, and “Δ G+AR1” showed the mean and standard deviation for the model’s cor(*g*_Rep1_,*g*_Rep2_) difference compared to the baseline G, G+2DSpl, and G+AR1 models, respectively. The “Δ G” and “Δ G+2DSpl” included the 2015, 2017, 2019, and 2020 field experiments; however, the 2015 field experiment could not be included for “Δ G+AR1” because of convergence issues. The (*) and (+) symbols denoted t-test p-values less than 0.05 and 0.1, respectively.

| Model | $\Delta$ G | $\Delta$ G+2DSpl | $\Delta$ G+AR1 |
| --- | --- | --- | --- |
| G+H3_RRID | 0.188 ± 0.094 (*) | 0.095 ± 0.045 (*) | 0.082 ± 0.053 (+) |
| G+H3_2DSplM | 0.123 ± 0.055 (*) | 0.029 ± 0.074 | 0.025 ± 0.09 |
| G+Havg_2DSplU | 0.12 ± 0.056 (*) | 0.026 ± 0.048 | 0.012 ± 0.057 |
| G+L_2DSplM | 0.065 ± 0.049 (*) | -0.028 ± 0.105 | -0.056 ± 0.123 |
| G+L_AR1U | 0.132 ± 0.077 (*) | 0.038 ± 0.035 (+) | 0.041 ± 0.031 (+) |
| G+L_RRID | 0.133 ± 0.041 (*) | 0.039 ± 0.043 (+) | 0.036 ± 0.049 |

Drawing from the simulation results in Figure 5, the 2DSplM and AR1M models may have had less overall accuracy due to the assumptions in Equation 6 and Equation 8, respectively, which restricted estimation of spatial covariance components between time points, and thereby negatively impacted the simulation when weaker correlations (< 0.9) across time points were used. In real data, as seen in Figure 2, Figure 3, Figure S8, Figure S9, and Figure S10, NLEE tended to be strongly correlated (>0.8) between time points; this may have explained the improvements seen in Table 5 when the 2DSplM NLEE were incorporated into second-stage genomic prediction. The RRAR1 and RR2DSpl models performed well in first-stage simulation; however, the second-stage genomic prediction was not particularly improved by these models potentially due to overfitting and a pronounced mismatch between the detected LEE and the causal effects. The RRID, AR1U, and 2DSplU models performed well in first-stage simulation and significantly improved the second-stage genomic prediction, indicating these models provided robust detection of spatial heterogeneity.

The second stage in the proposed approach could use soil data as an alternative to NLEE from the first-stage. Figure 7 illustrates differences in 2019_NYH2 against the baseline G (G Diff) for genomic heritability, model fit, and cor(*g*_rep1_, *g*_Rep2_) for GY, GM, and EH. The best eight models when modeling L (G+L) or FE (G+H) using soil data are illustrated in Figure 6; however, Figure S21 illustrates all of the models using soil data. Again, the baseline spatially corrected models, G+2DSpl and G+AR1, are shown. Included were the L models named G+L_RRSoilEC and G+L_RRSoilAlt, and the FE models named G+Favg_Soil_Alt, G+Favg_Soil_dAlt, G+Favg_Soil_d2Alt, G+Havg_Soil_EC, G+Havg_Soil_dEC, and G+Havg_Soil_d2EC. The soil information increased GM and EH heritability, and model fit for all traits, more than the baseline G+2DSpl and G+AR1 models; however, for all traits cor(*g*_rep1_, *g*_Rep2_) performed lower than the baseline G+2DSpl and G+AR1 models, particularly for GM and EH.

**Figure 7:**
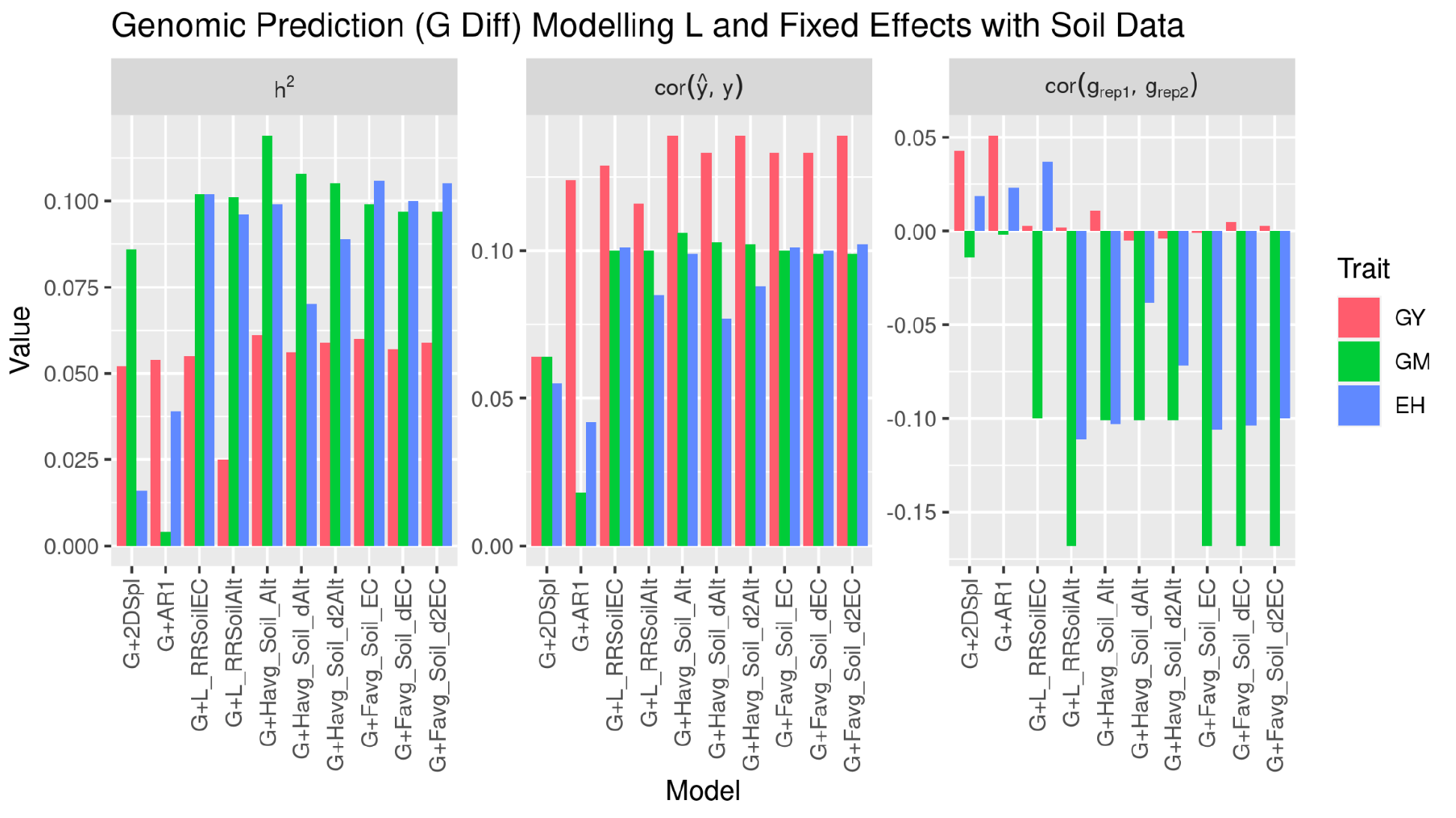
Differences compared to G (G Diff) for genomic heritability (*h*^2^), model fit (cor(*ŷ,y*)), and genotypic effect estimation across replicates (cor(*g*_rep1_, *g*_Rep2_)) in 2019_NYH2 for GY, GM, and EH, with soil data implemented as L (G+L) or as FE (G+H). The best eight models using soil altitude (Alt), soil electrical conductance (EC), and the first and second derivatives (dAlt, d2Alt, dEC, d2EC) were illustrated.

This approach to incorporate soil data did not improve GY cor(*g*_rep1_, *g*_Rep2_) however, this result was from a single field experiment in a single year. Similar to how the NDVI HTP itself did not correlate highly with GY while the spatial effects of NDVI correlated strongly with the spatial effects of GY, the spatial effects of the soil data may prove more beneficial for improving GY cor(*g*_rep1_, *g*_Rep2_) than the soil data itself. The soil data had much weaker correlations than the NDVI to agronomic traits, with a high of 0.07, 0.03, and 0.20 for GY, GM, and EH, respectively, compared to NDVI with a high of 0.67, 0.60, and 0.46 for GY, GM, and EH, respectively. Further indicating persistent spatial effects may be limiting the effectiveness of soil EC data in this study, Table 4 illustrates that including the soil data into RR models resulted in NLEE relatively well correlated with the soil elevation gradients, but negatively correlated with the soil EC data itself. The soil data were able to increase heritability; however, it may be a result of overfitting. Furthermore, soil data may need to be incorporated with weather information in order to effectively estimate the benefit or detriment of the local environmental effect. For instance, low elevation can be either beneficial or detrimental depending on rainfall.

## Conclusion

The proposed approach studied spatial heterogeneity in the field across time by answering the following two questions. Question 1: are NDVI LEE (NLEE), estimated by spatial effects and PE effects, consistently able to detect across the growing season the spatial heterogeneity affecting end-of-season agronomic traits? Question 2: can NLEE be used in the proposed two-stage models to improve spatial corrections for GP of agronomic traits?

Focusing on Question 1, in all years and for all agronomic traits, separating the random additive genetic effects resulted in stronger correlations between the agronomic trait spatial effects and NLEE than correlations between the agronomic traits and NDVI themselves. Furthermore, the NLEE from 2DSpl, AR1, and RR models consistently identified the same poorly performing regions in the field over the growing season, and identified substantially the same regions as the baseline GY and EH spatial effects.

Baseline GM spatial effects showed an inverted behavior with NLEE and were less localized than for GY and EH. The soil EC correlated most with NLEE from the 2DSplU and AR1U models across time. Therefore, spatial heterogeneity quantified by NLEE corresponded strongly with agronomic trait spatial effects and soil EC.

Focusing on Question 2, incorporating first-stage NLEE into the second-stage GP for GY, EH, and GM either as a covariance of random effects (L) or as fixed effects (FE), significantly improved heritability, model fit, and genotypic effect estimation across replicates. In simulation, the RRAR1 and RR2DSpl models performed strongly; however, only the RRID, AR1U, and 2DSplU models performed well in simulation and also improved the two-stage GP for agronomic traits. The RRID model made no spatial assumptions and performed consistently above average in simulation. The equilibrium between model generalizability and model over-specification when detecting NLEE was balanced most by the RRID, AR1U, and 2DSplU models.

Aerial image HTP provided greater understanding of spatial heterogeneity in the field, and when coupled into the proposed two-stage GP approach, enabled a more effective spatial correction than any of the baseline models (G+2DSpl, G+AR1, and M). Furthermore, the observed spatial heterogeneity could be partially explained using soil EC and elevation. Continued research into image features more informative than VI is needed. Additionally, further research is needed for the development of novel statistical approaches for integrating HTP across the growing season with end-of-season agronomic trait prediction. To these ends, larger datasets are required to evaluate the proposed approaches, and the continued aggregation of FAIR data is crucial.

## Data Availability

This study used phenotypic data of hybrid maize (*Zea mays* L.) field experiments part of the Genomes to Fields (G2F) program planted in 2015 (https://doi.org/10.25739/erxg-yn49), 2017 (https://doi.org/10.25739/w560-2114), 2019 (https://doi.org/10.25739/t651-yy97), and 2020 (https://doi.org/10.25739/hzzs-a865), named 2015_NYH2, 2017_NYH2, 2019_NYH2, and 2020_NYH2, respectively. The genotypic SNP marker data was also from the G2F program (https://doi.org/10.25739/frmv-wj25).

## Acknowledgements

The authors would like to thank the Genomes to Fields consortium for providing the NYH2 field experiments from 2015 to 2020 used in this study. This consortium involves more than 30 researchers representing more than 20 research institutions. Details about the initiative and publicly available resources can be found at www.Genomes2Fields.org. Thanks to Chris Hernandez, Peter Selby, Sam Bouabane, Ranjita Thapa, Simon Reinhard, Lynn Johnson, Seth Murray, Jacob Washburn, Filipe I. Matias, Annarita Marrano, and Felipe Sabadin for their help and suggestions on the image processing pipeline and the research more broadly.

## Supplemental Material

### File S1: Simulations

The first simulation process, the linear process, followed an evaluation of

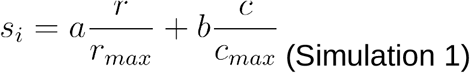

where *a* and *b* were randomly assigned numbers between 0 and 1, *r* and *c* were the current plot’s row and column position in the field, and *r*_*max*_ and *c*_*max*_ were the maximum row and column numbers in the field, respectively. The variable *s*_*i*_ was the simulated environmental effect for a specific experimental plot. Simulation 1 resulted in a constant gradient traversing the field, which increased as the row and column position increased. The second simulation process, the one-dimensional normal (1D-N) process, followed an evaluation of the univariate normal distribution equation below.

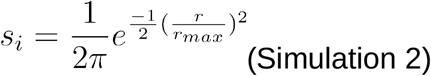

Simulation 2 resulted in a normal gradient which was constant across columns, but increased across the rows. The third simulation process, the two-dimensional normal (2D-N) process, followed an evaluation of the bivariate normal distribution equation

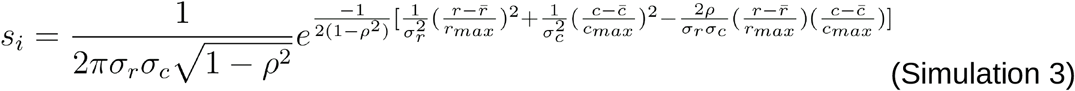

where *ρ* was a randomly assigned correlation between row number *r* and column number *c*, 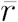 and 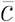 denoted the mean row and column numbers, respectively, *σ*_*r*_ and *σ*_*c*_ and denoted the standard deviations among the rows and columns, respectively. Simulation 3 resulted in a peak in the center of the field with normally distributed gradients decreasing across the rows and columns; the skew in the row and column gradients was controlled by *ρ*. The fourth simulation process, the separable autoregressive process (AR1xAR1), followed a multivariate normal distribution

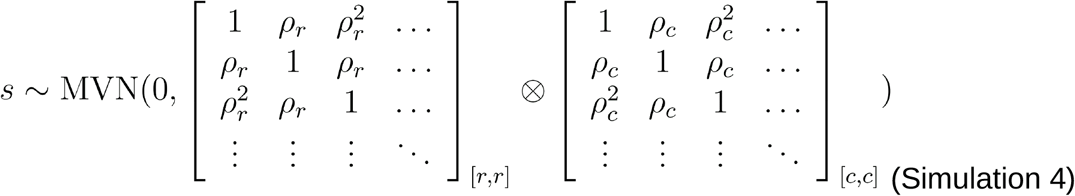

where *s* was a vector of simulated values, *ρ*_*c*_ and *ρ*_*r*_ and were randomly assigned correlations among the rows and columns, respectively. Simulation 4 explicitly defined a decay in the correlation between experimental plots based on their proximity on a unit-by-unit basis (e.g. *ρ* > *ρ*^2^ > *ρ*^*3*^ >…). The multivariate normal distribution was implemented using the MASS package (Venables and Ripley 2002). The fifth simulation process, the random process, followed

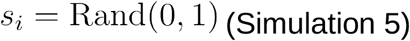

and produced random values between 0 and 1. Simulation 5 was used as a control to determine the effect of random noise. The sixth simulation process, the real data (RD) process, was intended to use the collected soil EC data directly

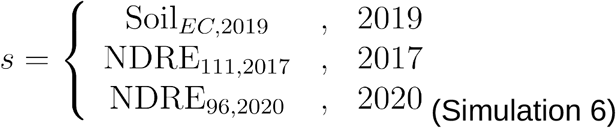

where the vector of simulated values was the soil EC data from 2019. In 2017 and 2020 soil data was not available, so NDRE at 111 and 96 days after planting, respectively, was used instead. It was important to use a measurement with no missing data, therefore, the HTP measurement of NDRE was suitable.

For each of the six simulations described, three approaches were explored for generating the simulated field effect across time. The first approach defined the simulated field effect as constant through time. This can be written as 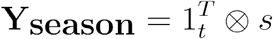, where Y_season_ was a matrix representing all simulated values across the season, 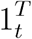 was a transposed vector of ones with length equal to the number of time points *t*, and *s* contained the simulated values from one of the six aforementioned simulation processes. Figure S5 illustrated heatmaps for all six simulation processes, constant over 12 timepoints. The second and third approaches generated simulated values which were 90% and 75%, respectively, correlated through time. This was accomplished by specifying a correlation structure C between time points as 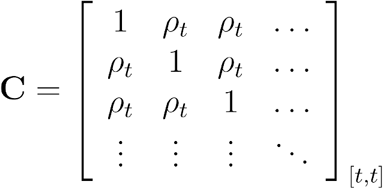, where *ρ*_*t*_ was the correlation between time points, set to either 0.75 or 0.90, and the dimensions of were [*t,t*] to denote the number of time points. Simulated correlated values Y_season_ were generated across the growing season by first taking the Cholesky decomposition, D, of *C* as C = D^T^D. Then, 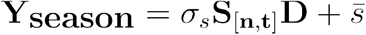 was defined, where *σ*_*s*_ and 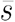 are the standard deviation and mean of the target simulated values *s*, respectively. The matrix S[n,t] contained the normalized simulated values *s* in the first column, followed by columns initialized by a unit normal distribution resulting in *t* columns. Figure S6 illustrated heatmaps for all six simulation processes over 12 time points which were 90% correlated to each other.

### Supplemental Figures

**Figure S1:**
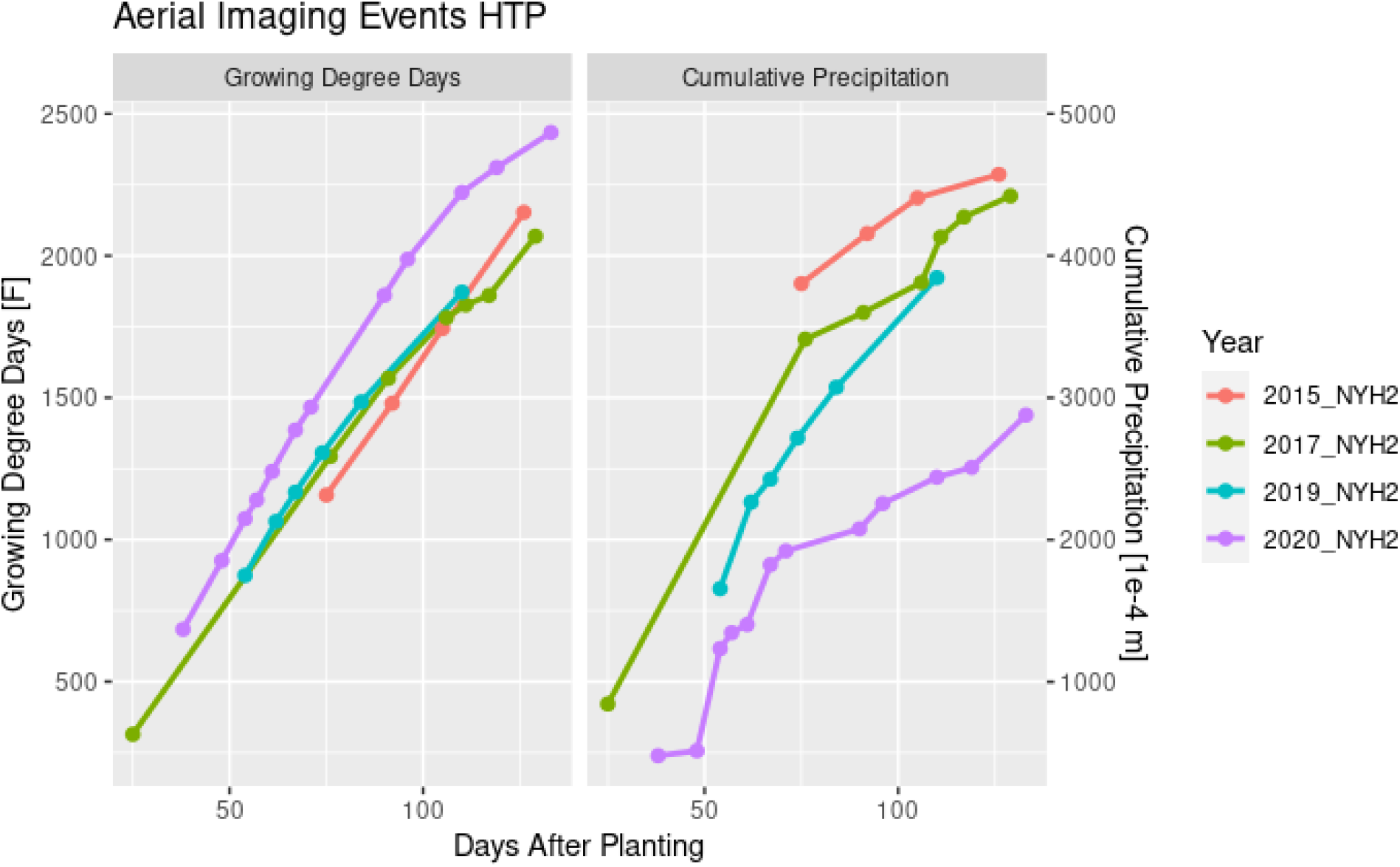
Aerial imaging events with HTP extracted for the hybrid maize experiments in 2015, 2017, 2019, and 2020. Growing degree days (GDD) and cumulative precipitation (CP) were plotted against the days after planting (DAP) of the imaging events. All field experiments were located in Musgrave Research Station, though planted in distinct fields, and weather data was sourced for the GHCND:USC00300331 ground station via the NOAA NCEI NCDC database.

**Figure S2:**
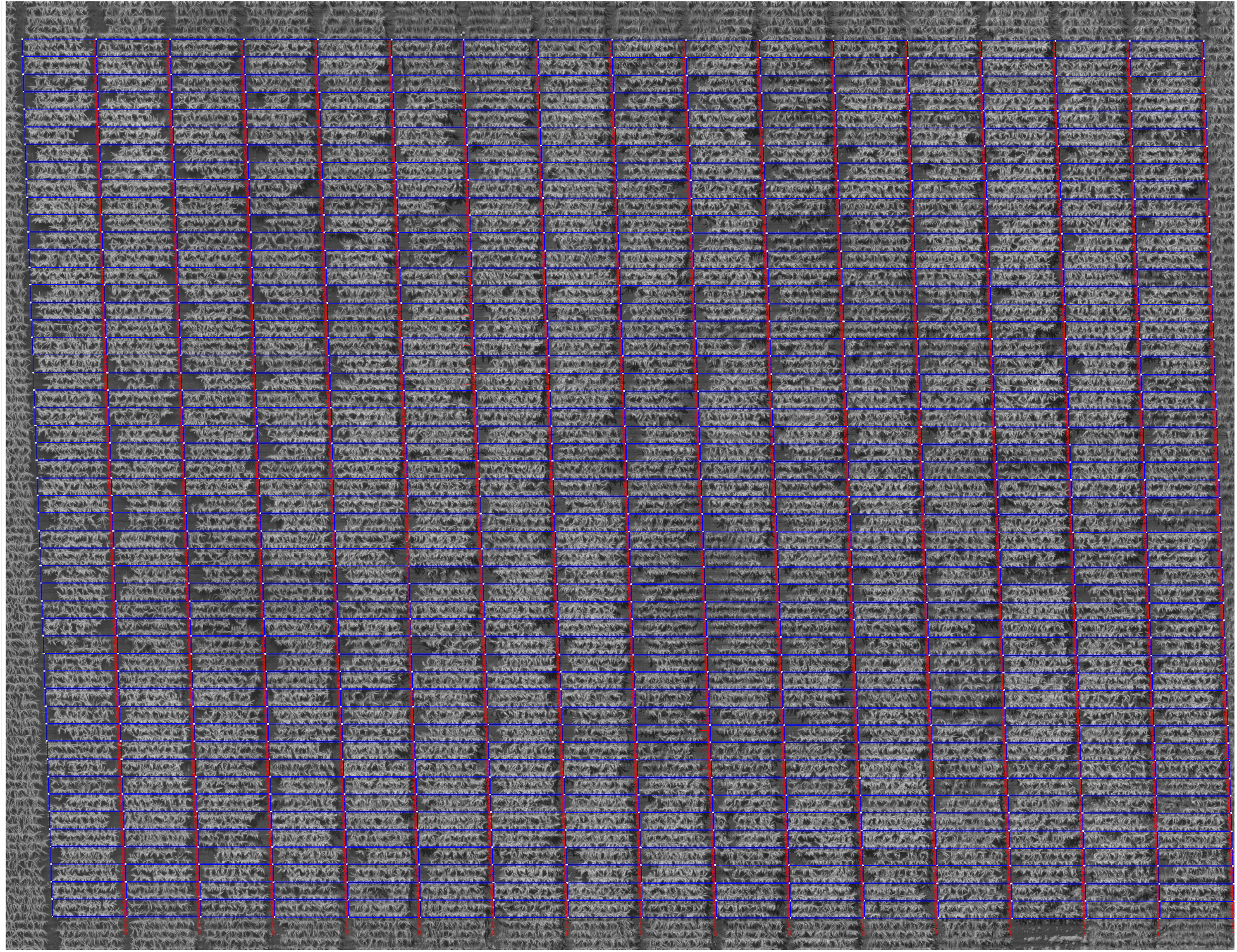
Illustrated is a representative reflectance orthophotomosaic raster image. This image was of 2019_NYH2 on Aug 15, 2019 and was in the near-infrared (NIR) spectra. The original image had dimensions of 6488 by 4987 pixels allowing about 1cm per pixel resolution and was available from https://imagebreed.org/data/images/image_files/26/05/f9/3e/ab9b340016a1db73f9c743cf/imagegoCo.png. The overlaid blue polygons represented plot-polygons drawn in ImageBreed to segment plot-images for high-throughput phenotype (HTP) extraction.

**Figure S3:**
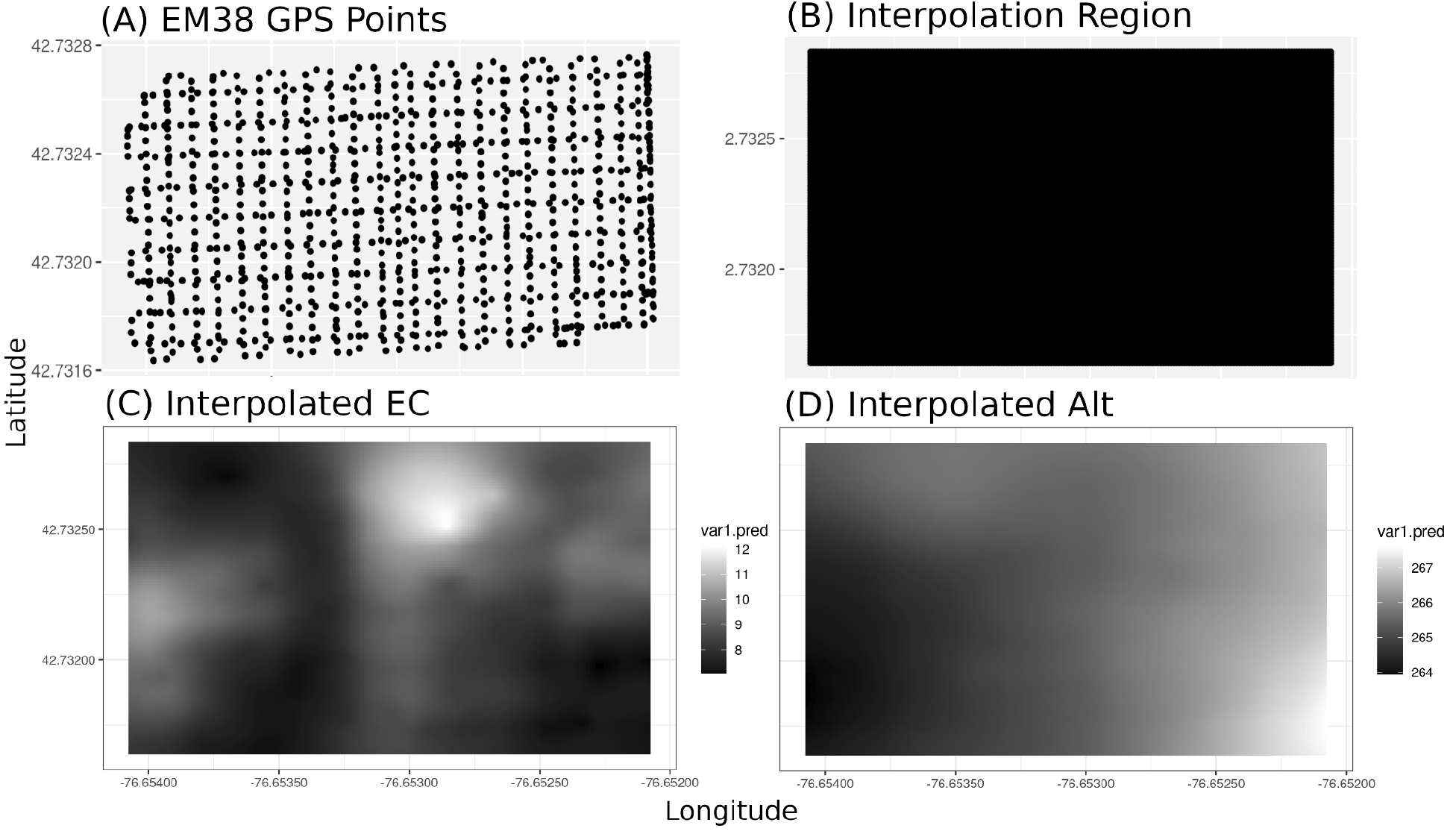
Illustrated are the 2019_NYH2 soil survey interpolation maps. (A) illustrated the raw EM38 soil survey GPS data collected using a dual-serpentine path. (B) illustrated the region to interpolate into with a 0.00001 WGS84 resolution across 200 by 120 cells. Finally, (C) and (D) showed the soil EC and altitude, respectively, interpolated across the field using ordinary Kriging.

**Figure S4:**
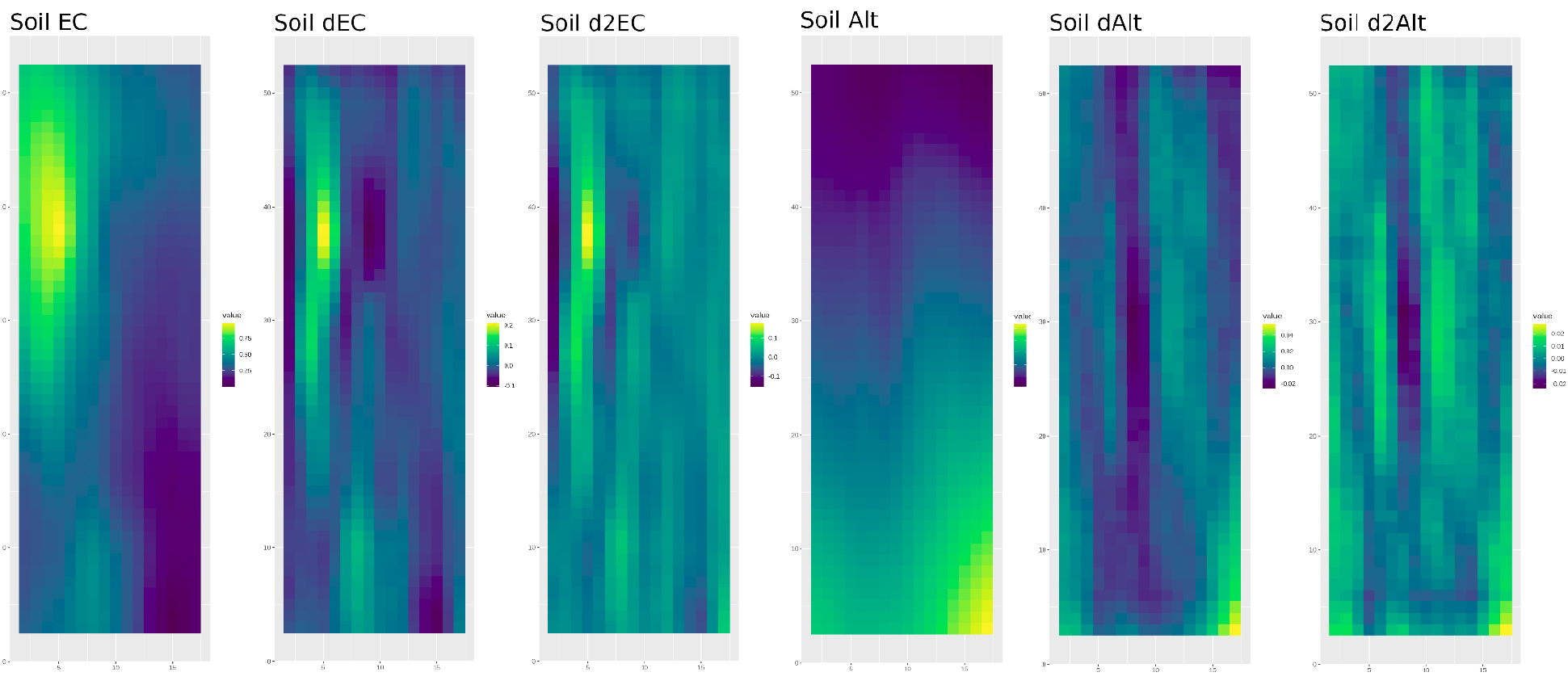
Soil EC and altitude was extracted for the experimental plots in 2019_NYH2. First and second two-dimensional numerical derivatives were computed. The heatmaps illustrated are of these soil measurements for the 800 experimental plots across 16 columns and 50 rows.

**Figure S5:**
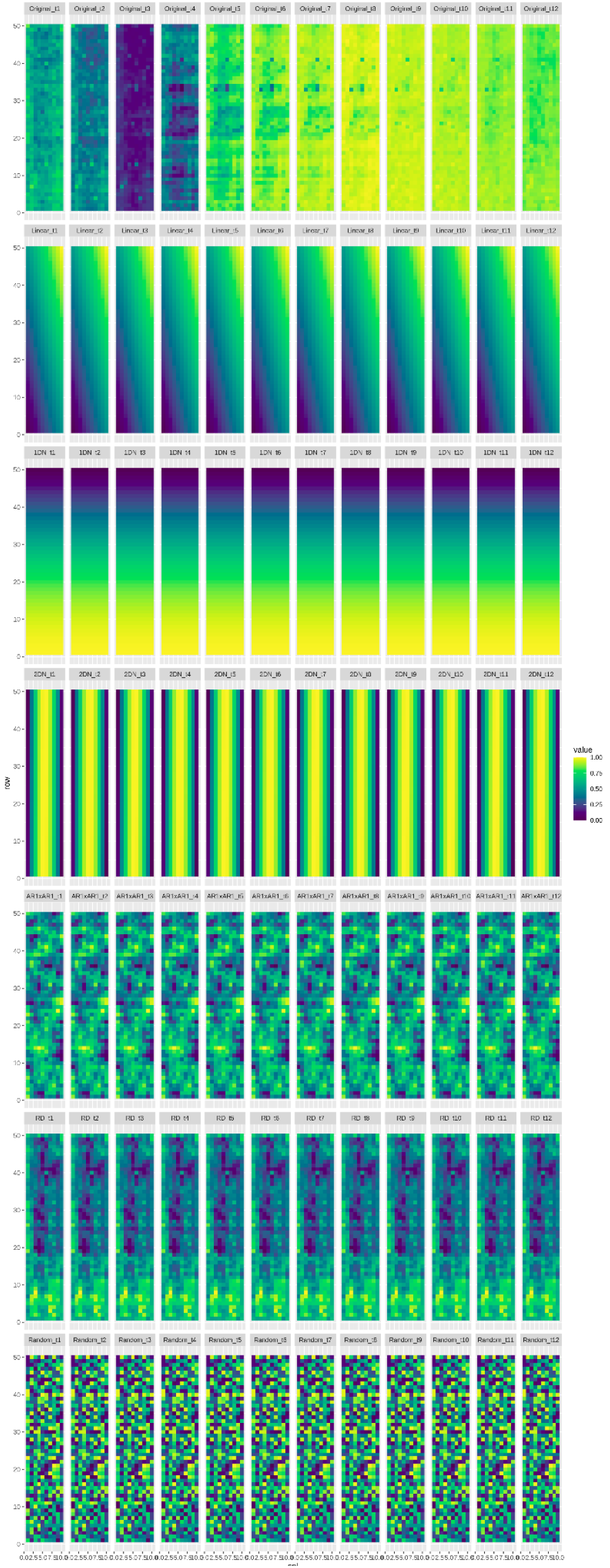
Illustrated are example heatmaps of the six simulation processes, constant through time. At the top, the actual 2020_NYH2 NDVI phenotype across the 12 imaging events was shown, followed by: the linear simulation (Simulation 1), the 1D-N (Simulation 2), the 2D-N (Simulation 3), the separable autoregressive (Simulation 4), the random (Simulation 5), and the real data (Simulation 6) processes, sequentially.

**Figure S6:**
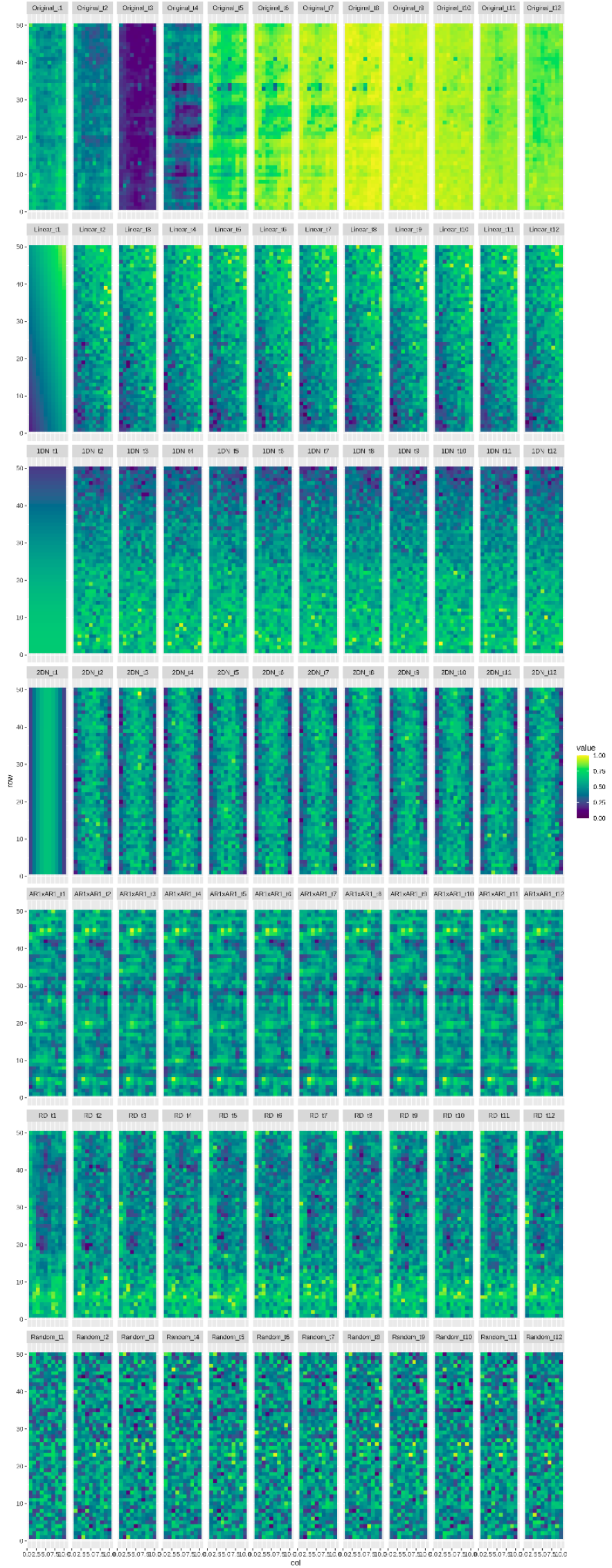
Illustrated are example heatmaps of the six simulation processes, 90% correlated across time. At the top, the 2020_NYH2 NDVI phenotype across the 12 imaging events was shown, followed by: the linear simulation (Simulation 1), the 1D-N (Simulation 2), the 2D-N (Simulation 3), the separable autoregressive (Simulation 4), the random (Simulation 5), and the real data (Simulation 6) processes, sequentially.

**Figure S7:**
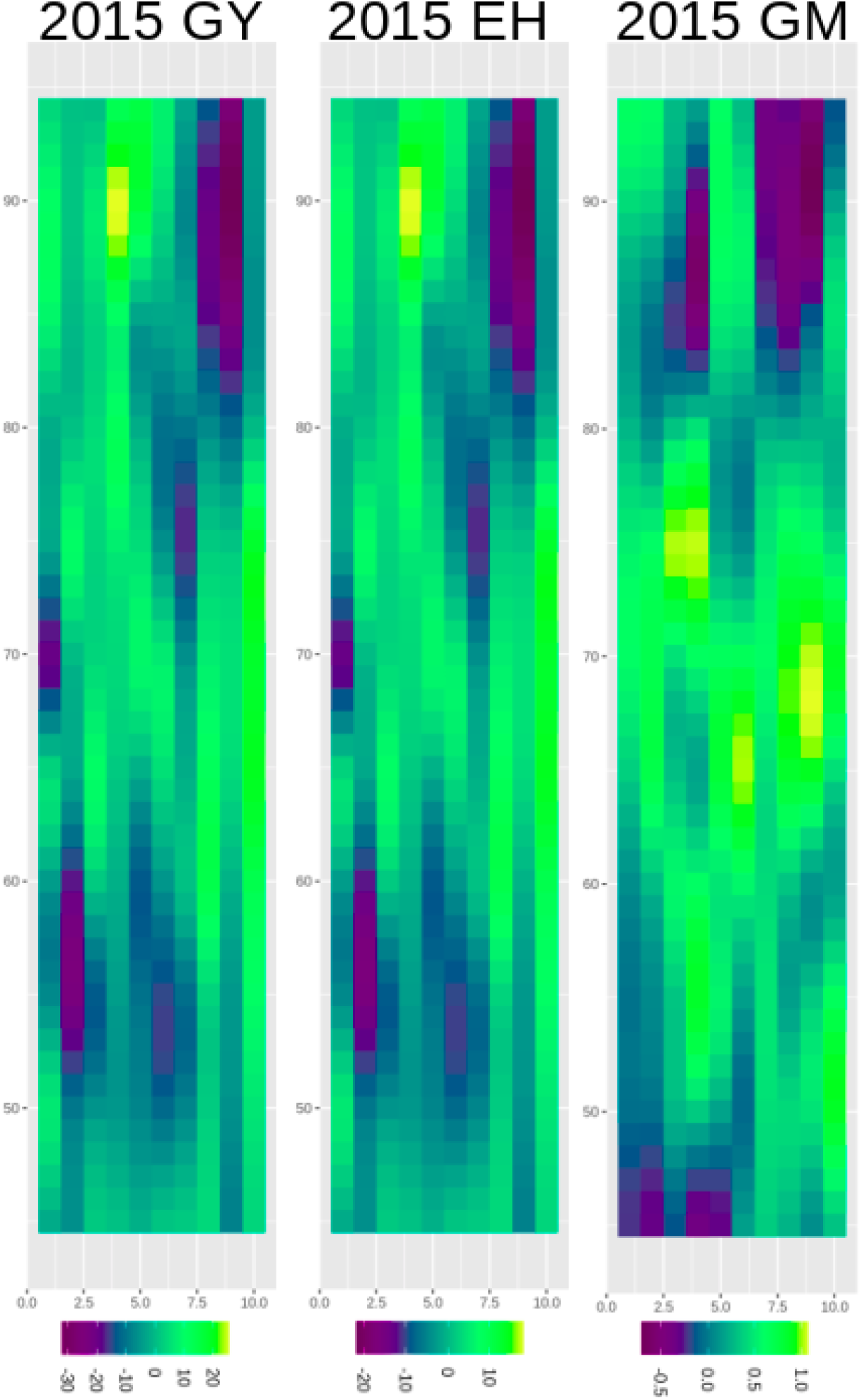
Illustrated are the two-dimensional spline (2DSpl) spatial effects for grain yield (GY), grain moisture (GM), and ear height (EH) in the 2015_NYH2 field experiment. Spatial effects are drawn over the 500 experimental plots across 10 rows and 50 columns.

**Figure S8:**
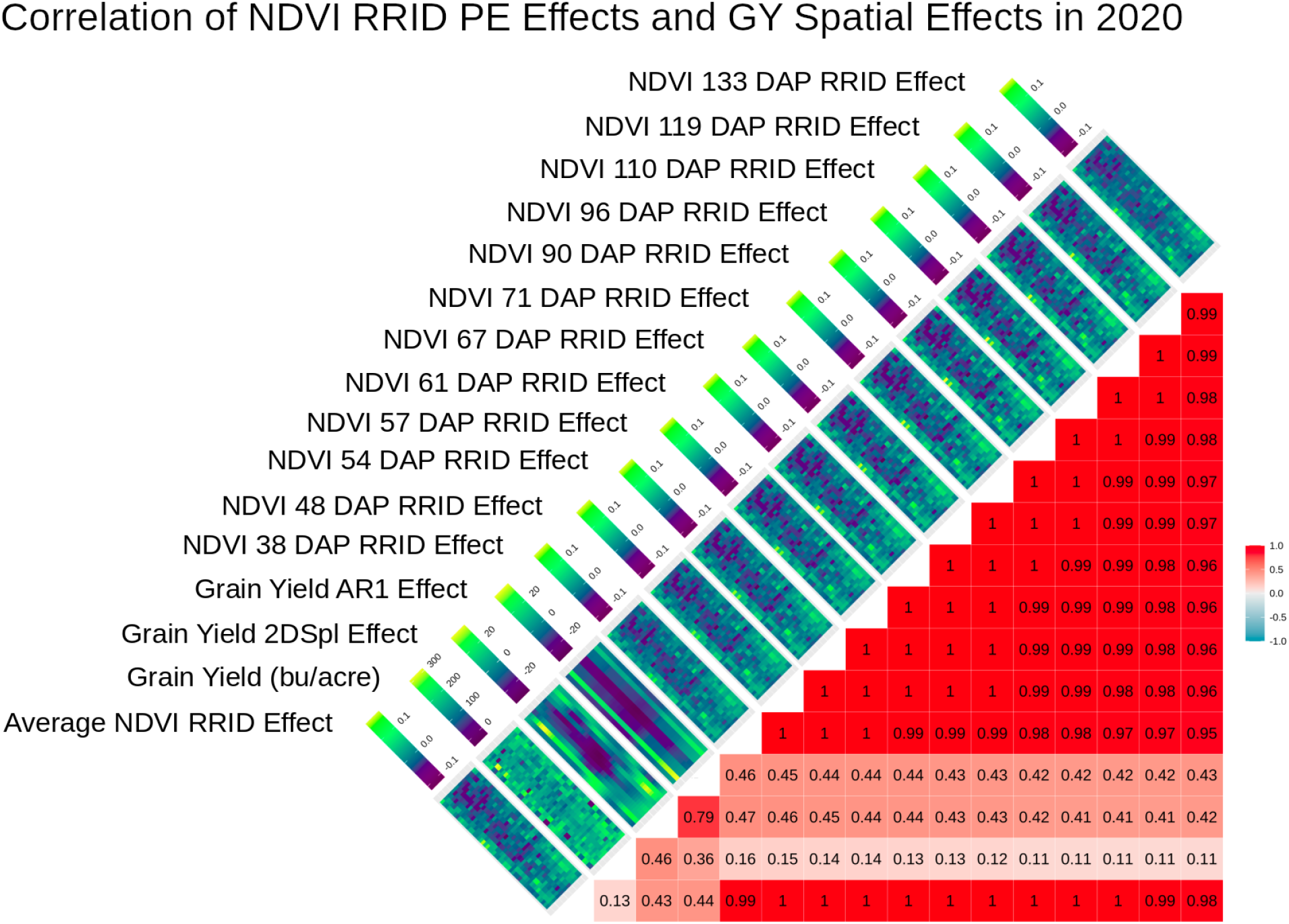
The NDVI RRID PE effects for 12 time points across the 2020_NYH2 growing season correlated with grain yield (GY) and the 2DSpl and AR1 spatial effects of GY. The NDVI PE and the GY spatial effects correlate at a value of 0.5 at 38 days after planting (DAP), and correlate above 0.4 across the growing season.

**Figure S9:**
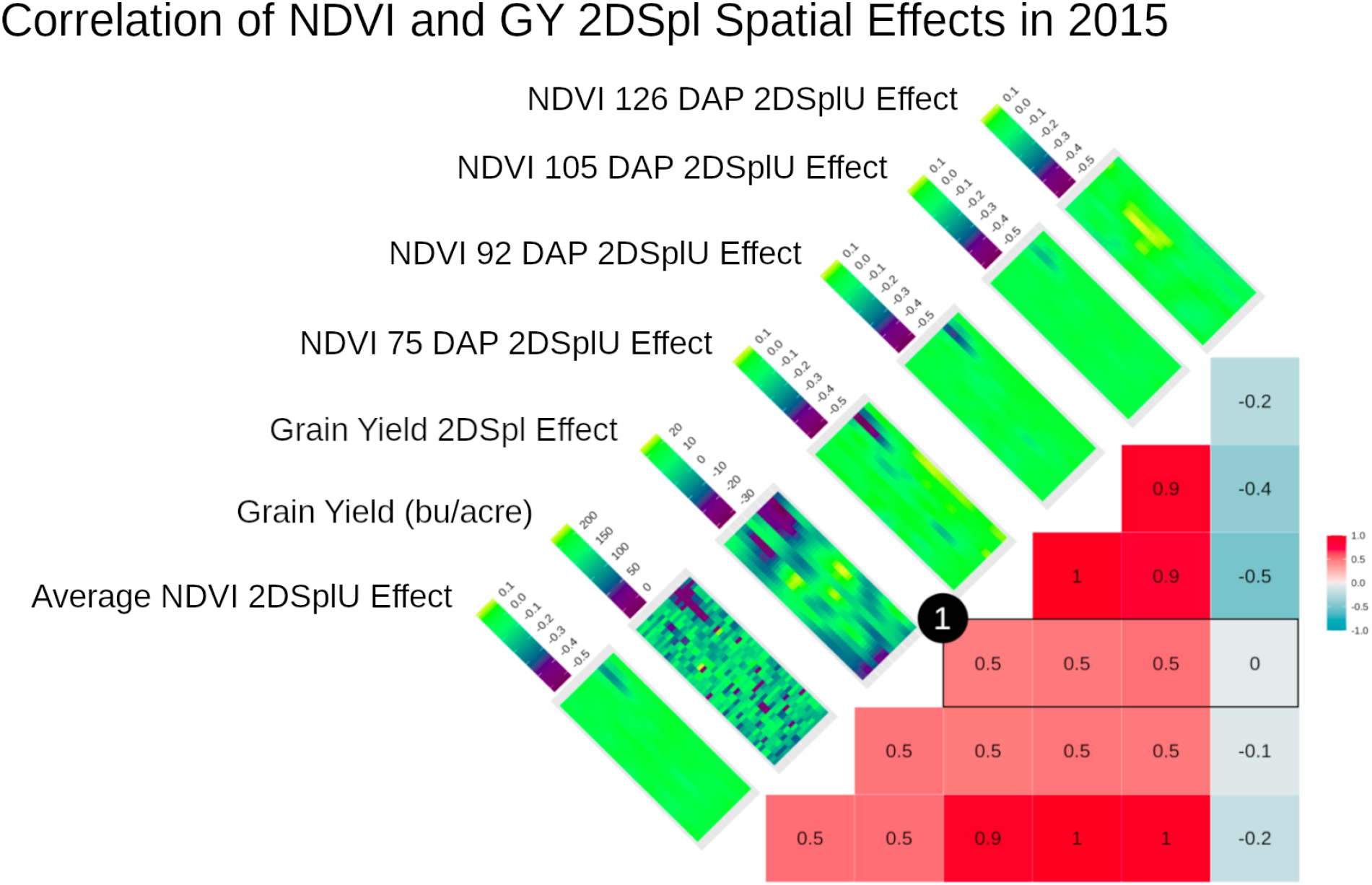
Shown are the two-dimensional spline (2DSpl) spatial effects for grain yield (GY) correlated with NDVI univariate 2DSpl (2DSplU) local environmental effects (LEE) for 4 time points in the 2015_NYH2 experiment. As demonstrated in (1), correlations of 0.5 were observed at 75, 92, and 105 days after planting (DAP). The corresponding heatmaps drew the values over the rows and columns of the experimental plots, and consistently identified one poorly performing region in the field.

**Figure S10:**
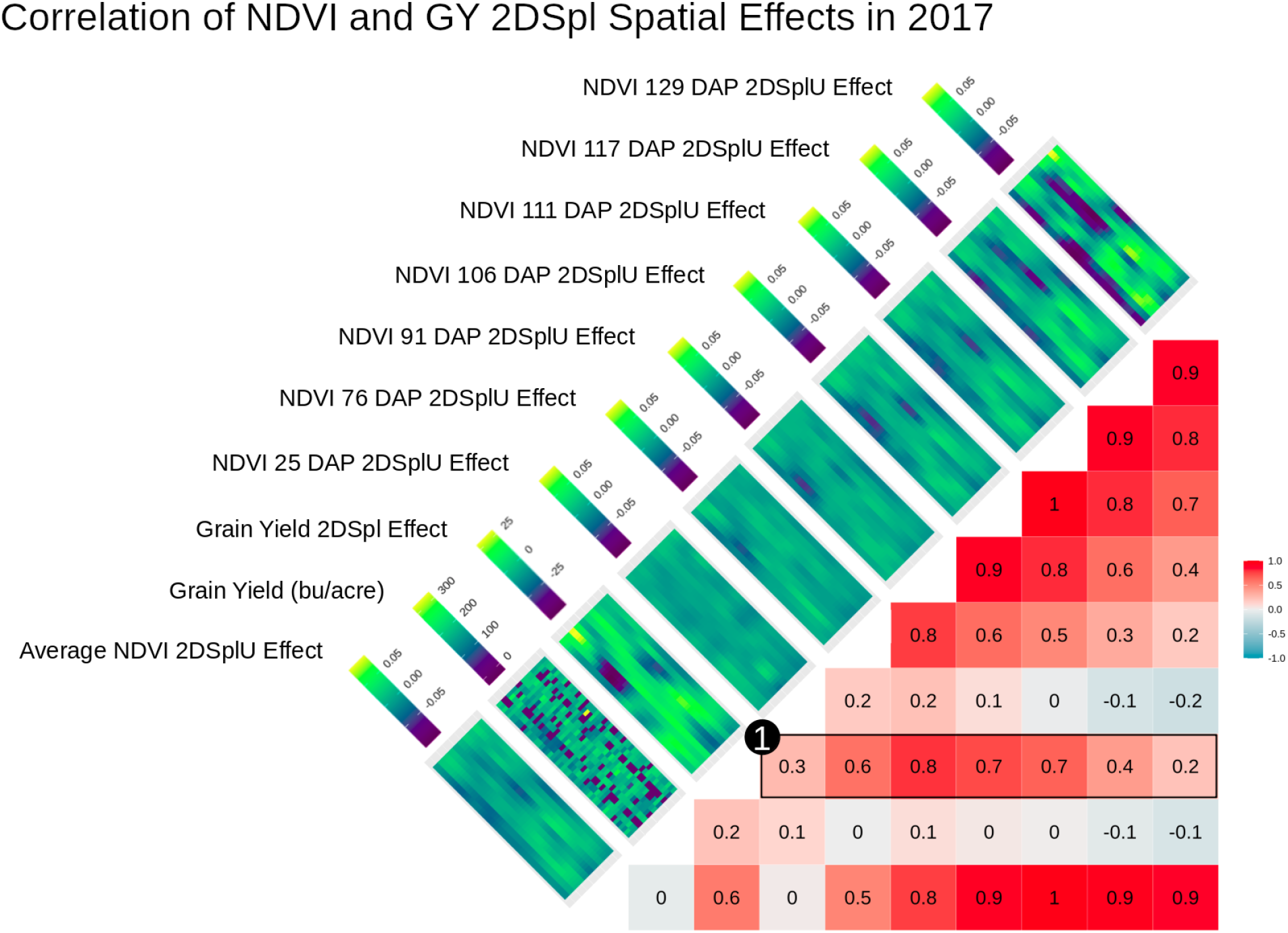
Shown are the two-dimensional spline (2DSpl) spatial effects for grain yield (GY) correlated with NDVI univariate 2DSpl (2DSplU) local environmental effects (LEE) for 7 time points in the 2017_NYH2 experiment. As demonstrated in (1), correlations of 0.2 to 0.8 were observed with the highest correlation at 91 days after planting (DAP). The corresponding heatmaps drew the values over the rows and columns of the experimental plots, and consistently identified three distinct poorly performing regions in the field.

**Figure S11:**
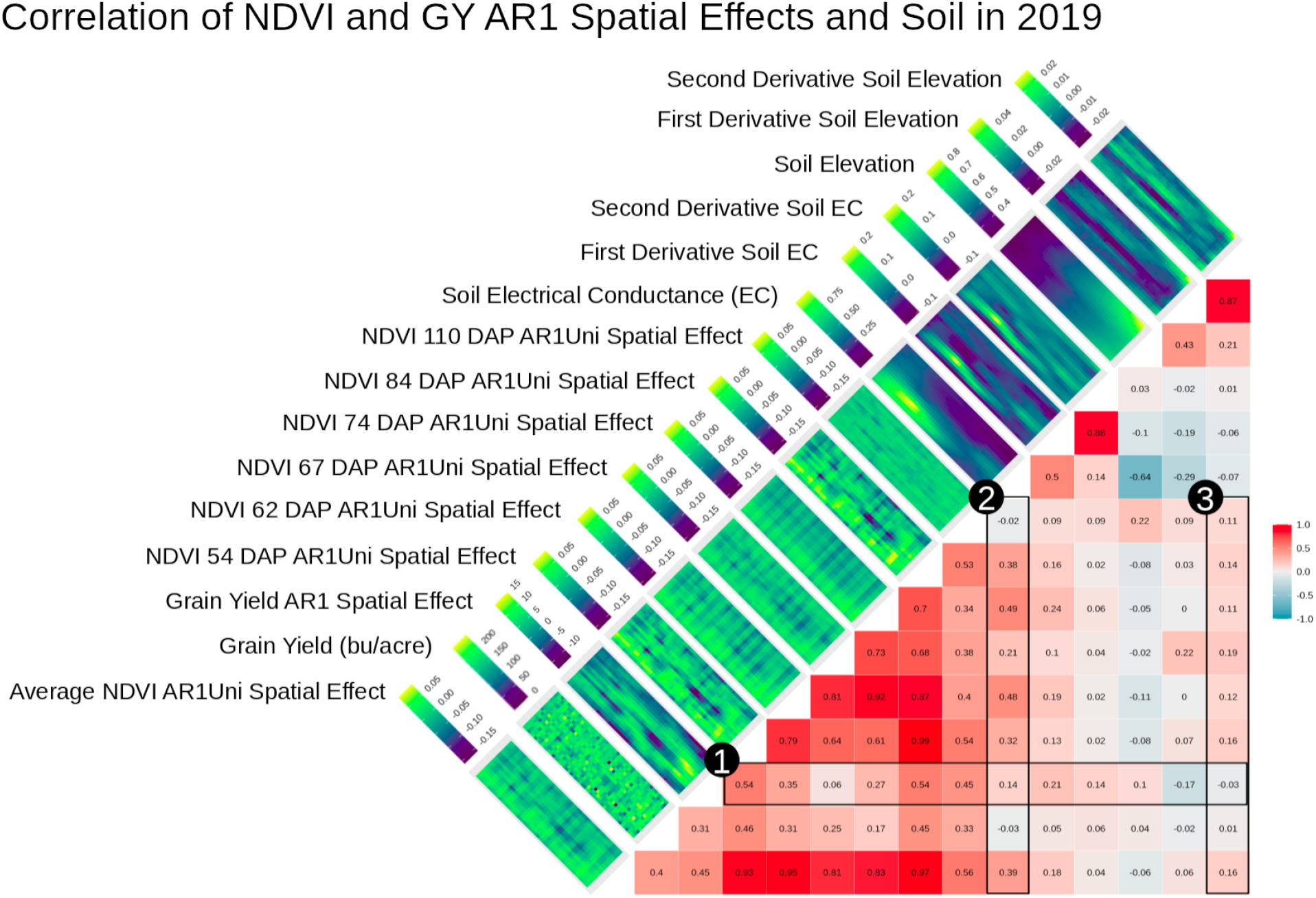
Drawn are the 2019_NYH2 AR1U NLEE observed over 6 time points correlated with GY and AR1 spatial effects of GY. Corresponding heatmaps showed values over the rows and columns of all experimental plots in the field, revealing similar spatial patterns. As indicated by (1), correlations of 0.1 to 0.5 between GY AR1 and the AR1U NLEE were found throughout the growing season. The highest correlation of 0.5 was observed at 110 DAP and 54 DAP. Soil EC and Alt, as well as the first and second numerical two-dimensional derivatives, were included with (2) showing correlations up to 0.5 for EC and (3) showing correlations up to 0.2 for d2Alt. The analog in Figure 3 showed that the 2DSplU NLEE were more highly correlated with the soil parameters than the AR1U NLEE, and the 2DSplU NLEE were more highly correlated with GY 2DSpl effects than the AR1U NLEE were correlated with the GY AR1 effects.

**Figure S12:**
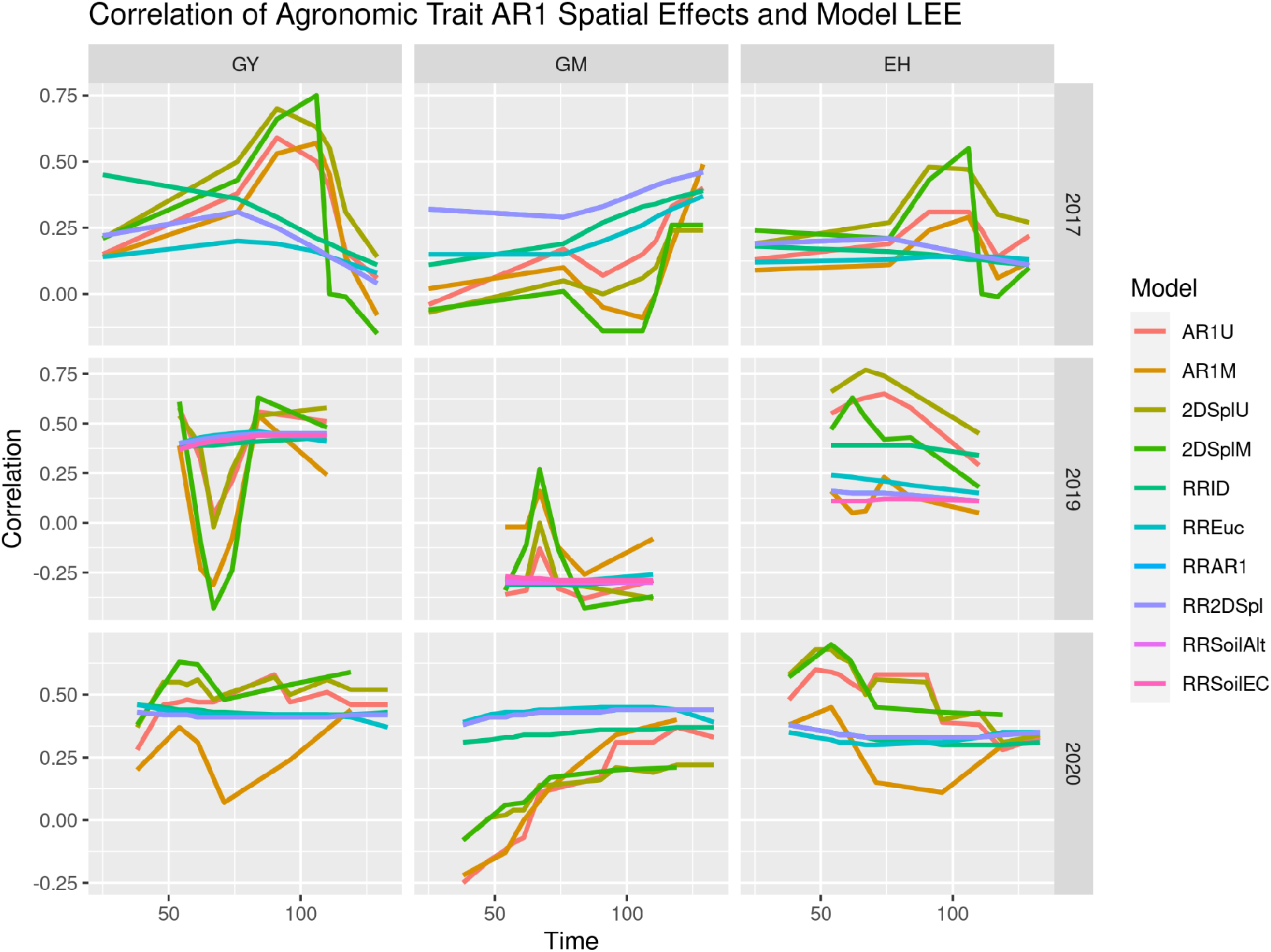
Illustrated are correlations between AR1 spatial effects of agronomic traits and NLEE from various models. Spatial effects for GY, GM, and EH in the 2017_NYH2, 2019_NYH2, and 2020_NYH2 field experiments were compared against NLEE across the growing season. Models run on traits in a given year showed similarities, with GY and EH following tandem trends, and GY and GM showing inverted patterns. 2015_NYH2 not included because convergence was not possible. The 2DSpl analog in Figure 4 illustrated similar patterns.

**Figure S13:**
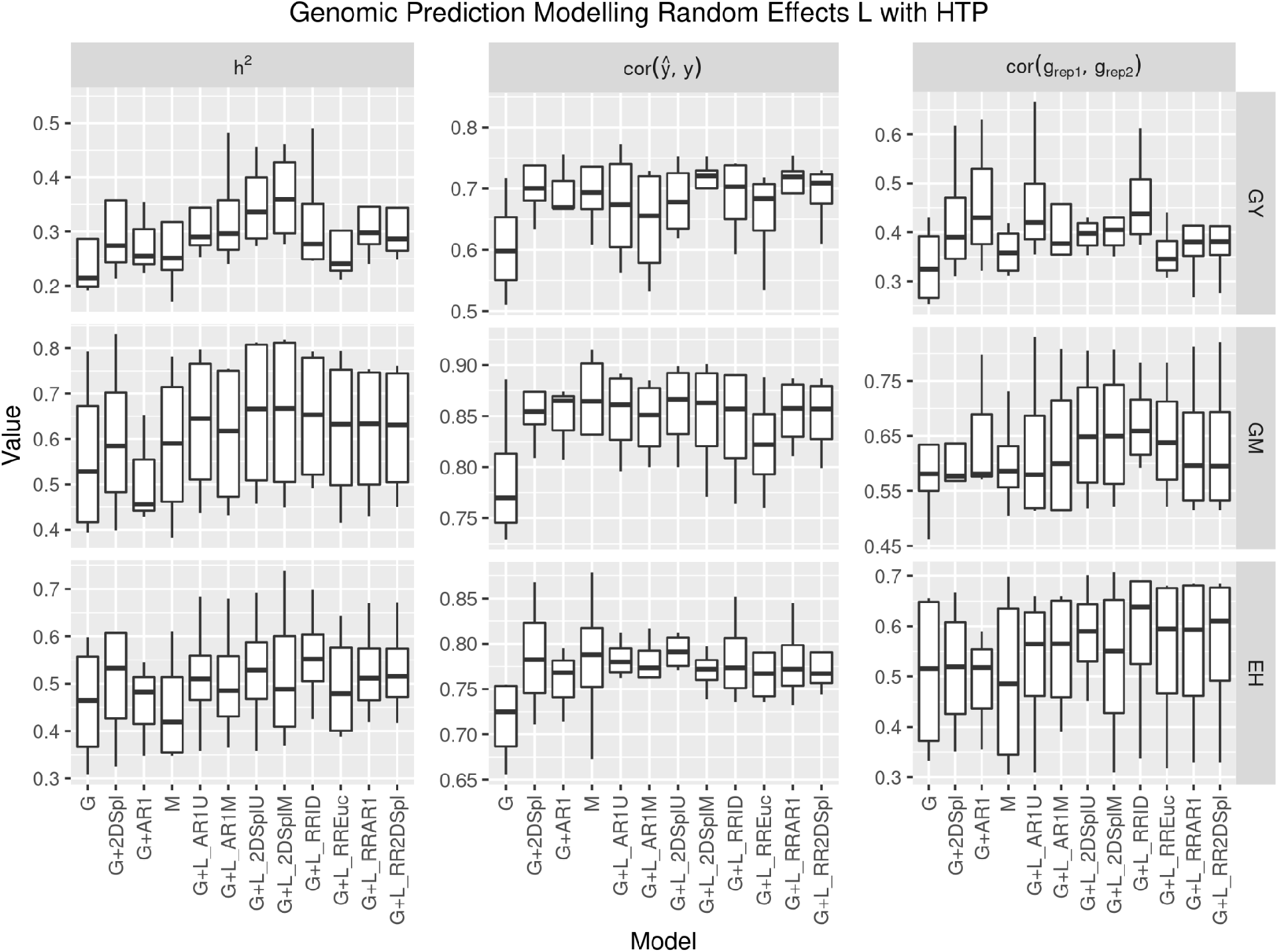
Genomic heritability (*h*^2^), model fit (cor(*ŷ,y*)), and genotypic effect estimation across replicates (cor(*g*_rep1_, *g*_Rep2_)) in the four years for GY, GM, and EH, with NLEE implemented as L. The models G, G+2DSpl, and G+AR1 were baseline GBLUP and spatially corrected models, respectively, and M was a baseline multi-trait model. Models have L defined using NLEE of corresponding name.

**Figure S14:**
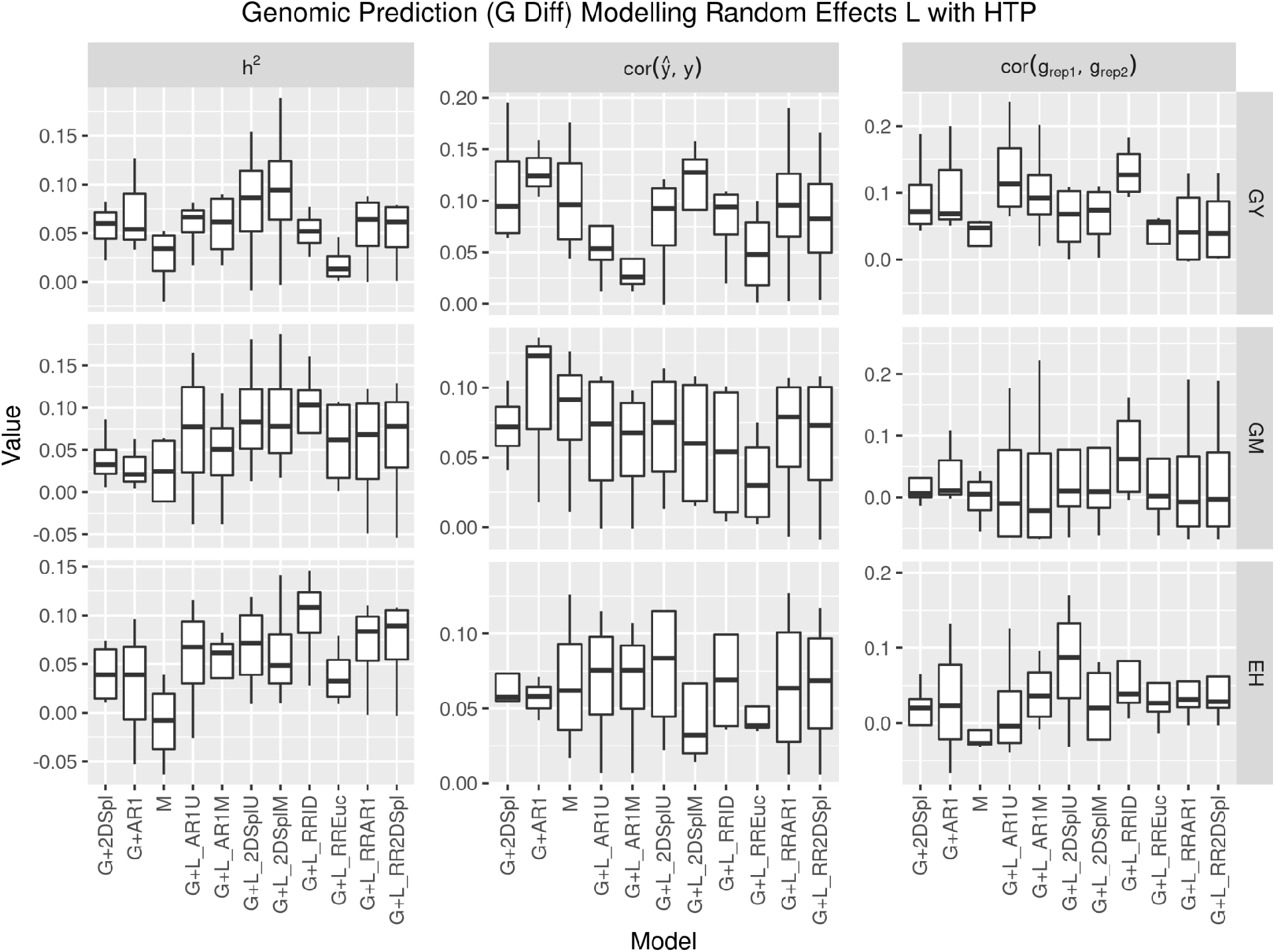
The difference in model genomic heritability (*h*^2^), model fit (cor(*ŷ,y*)), and genotypic effect estimation across replicates (cor(*g*_rep1_, *g*_Rep2_))with G (G Diff) in the four years for GY, GM, and EH, with NLEE implemented as L. The models G+2DSpl and G+AR1 were baseline spatially corrected GBLUP models, respectively, and M was a baseline multi-trait model. Models have L defined using NLEE of corresponding name.

**Figure S15:**
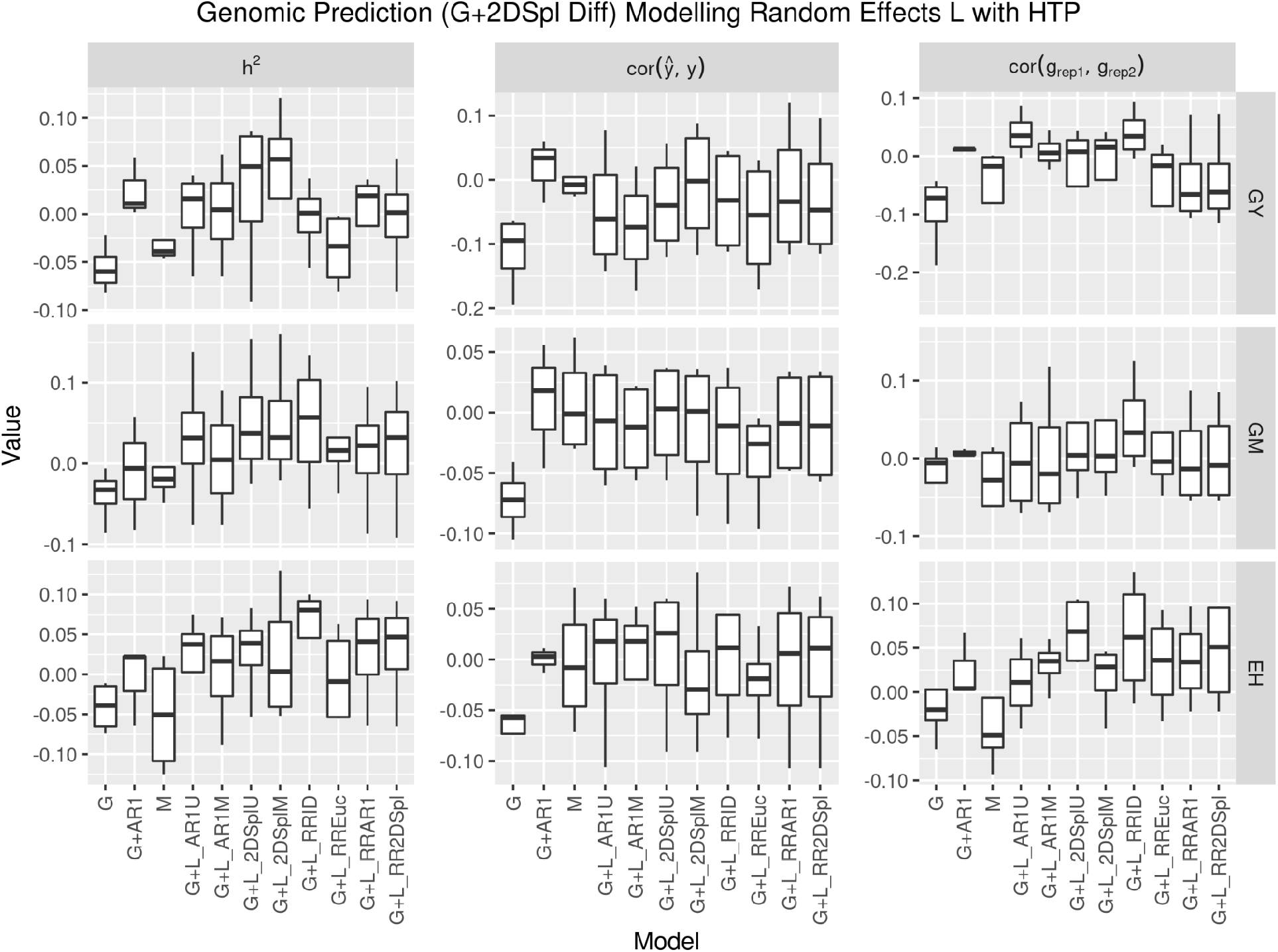
The difference in model genomic heritability (*h*^2^), model fit (cor(*ŷ,y*)), and genotypic effect estimation across replicates (cor(*g*_rep1_, *g*_Rep2_))with 2DSpl spatially corrected G (G+2DSpl Diff) in the four years for GY, GM, and EH, with NLEE implemented as L. The models G and G+AR1 were baseline GBLUP and spatially corrected models, respectively, and M was a baseline multi-trait model. Models have L defined using NLEE of corresponding name.

**Figure S16:**
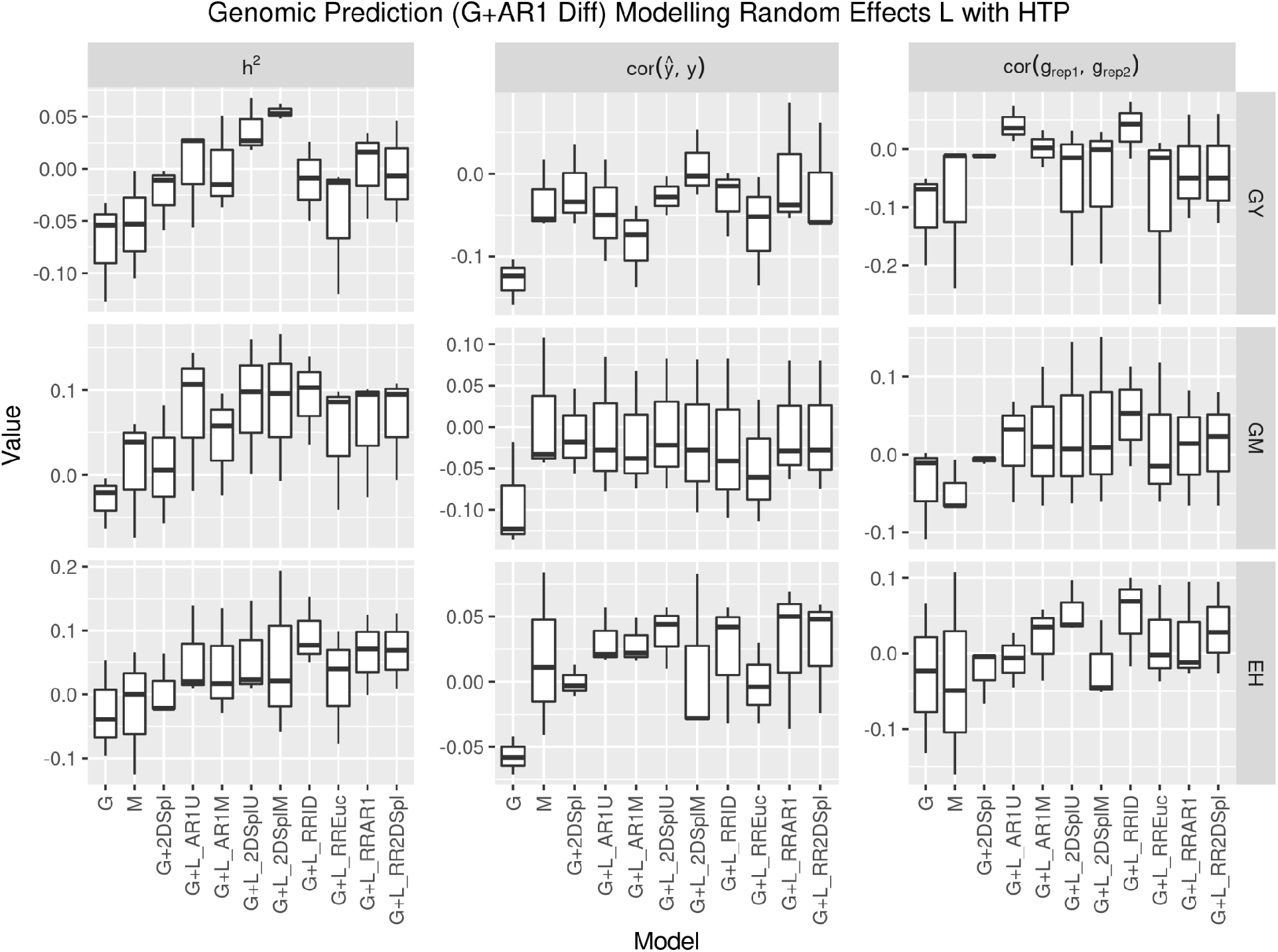
The difference in model genomic heritability (*h*^2^), model fit (cor(*ŷ,y*)), and genotypic effect estimation across replicates (cor(*g*_rep1_, *g*_Rep2_))with AR1 spatially corrected G (G+AR1 Diff) in the four years for GY, GM, and EH, with NLEE implemented as L. The models G and G+2DSpl were baseline GBLUP and spatially corrected models, respectively, and M was a multi-trait model. Models have L defined using NLEE of corresponding name.

**Figure S17:**
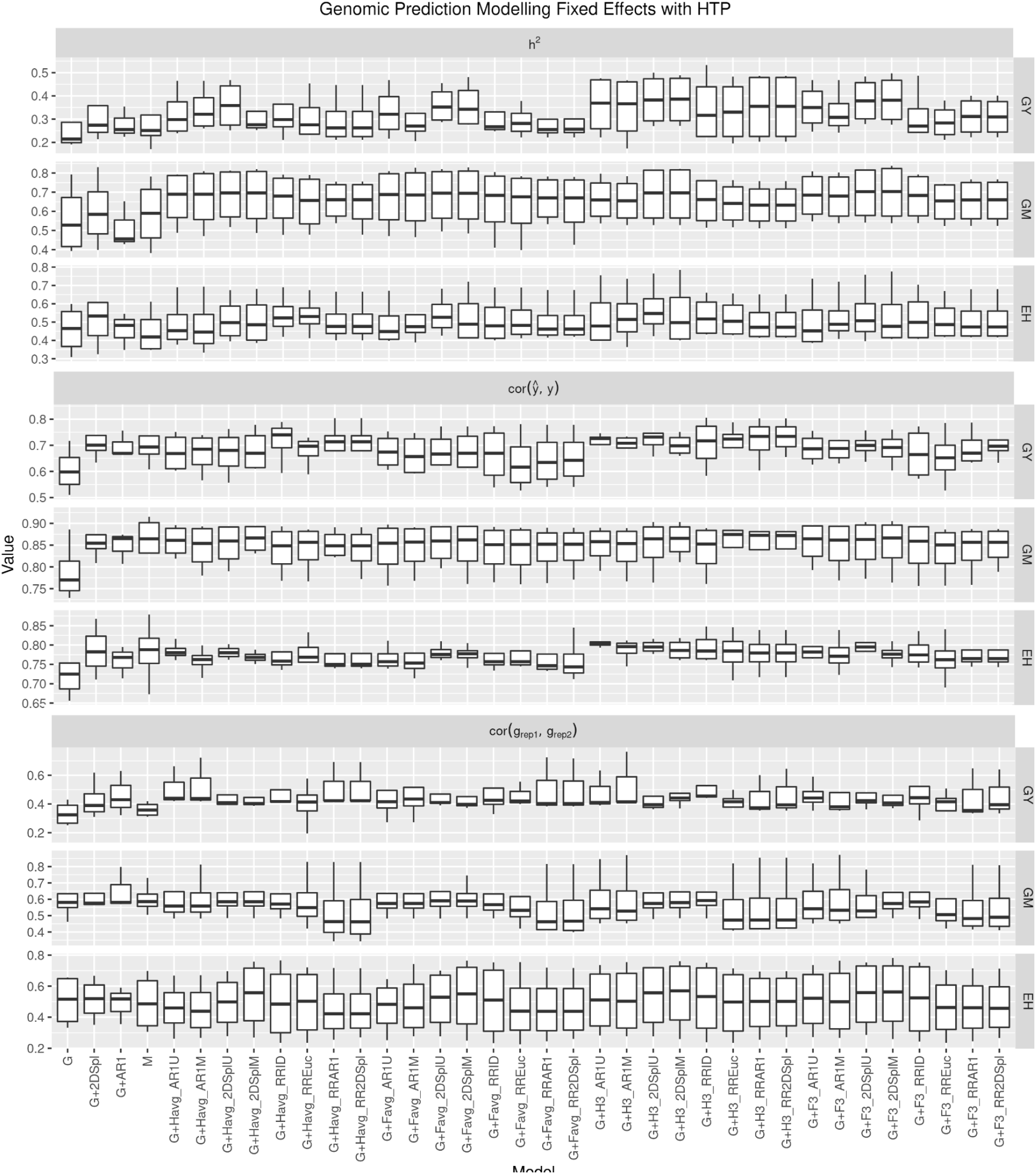
Genomic heritability(*h*^2^), model fit(cor(*ŷ,y*)), and genotypic effect estimation across replicates (cor(*g*_rep1_, *g*_Rep2_)) in the four years for GY, GM, and EH, with NLEE implemented as FE. The models G, G+2DSpl, and G+AR1 were baseline GBLUP and spatially corrected models, respectively, and M was a baseline multi-trait model. Models have FE defined using NLEE of corresponding name.

**Figure S18:**
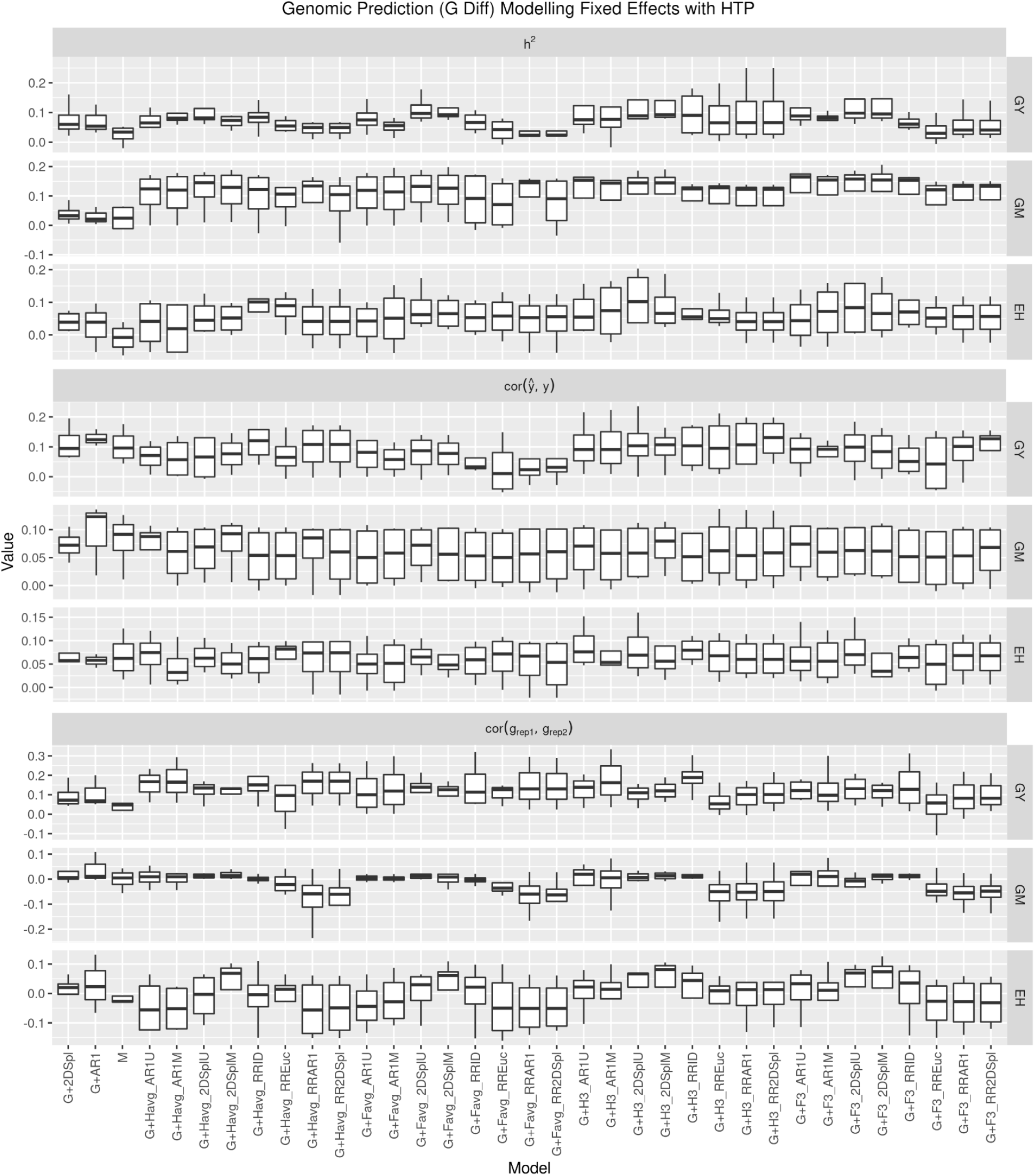
The difference in model genomic heritability (*h*^2^), model fit(cor(*ŷ,y*)), and genotypic effect estimation across replicates (cor(*g*_rep1_, *g*_Rep2_))with G (G Diff) in the four years for GY, GM, and EH, with NLEE implemented as FE. The models G+2DSpl and G+AR1 were baseline spatially corrected GBLUP models, respectively, and M was a baseline multi-trait model. Models have FE defined using NLEE of corresponding name.

**Figure S19:**
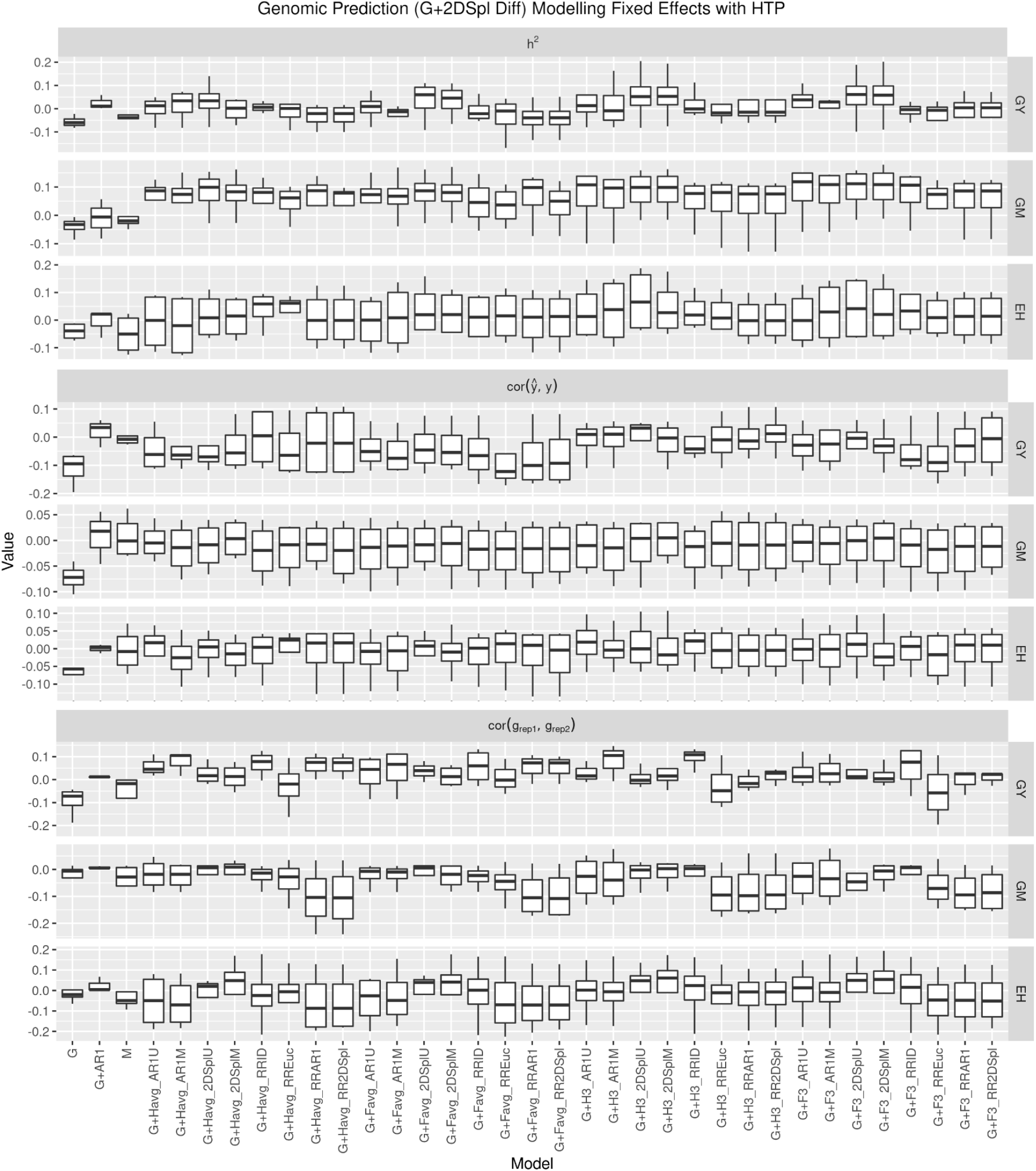
The difference in model genomic heritability (*h*^2^), model fit (cor(*ŷ,y*)), and genotypic effect estimation across replicates (cor(*g*_rep1_, *g*_Rep2_))with 2DSpl spatially corrected G (G+2DSpl Diff) in the four years for GY, GM, and EH, with NLEE implemented as FE. The models G and G+AR1 were baseline GBLUP and spatially corrected models, respectively, and M was a baseline multi-trait model. Models have FE defined using NLEE of corresponding name.

**Figure S20:**
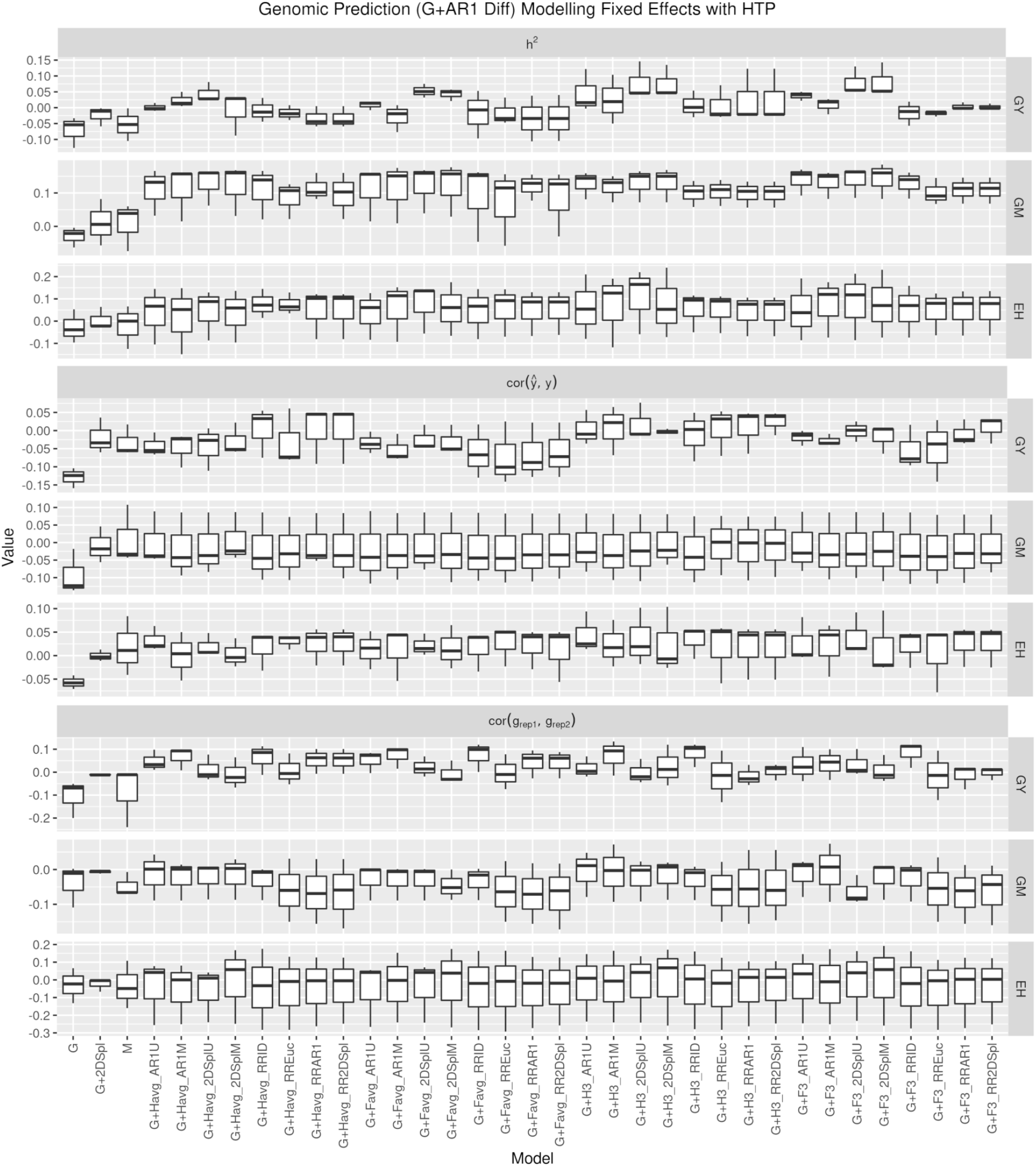
The difference in model genomic heritability (*h*^2^), model fit(cor(*ŷ,y*)), and genotypic effect estimation across replicates (cor(*g*_rep1_, *g*_Rep2_))with AR1 spatially corrected G (G+AR1 Diff) in the four years for GY, GM, and EH, with NLEE implemented as FE. The models G and G+2DSpl were baseline GBLUP and spatially corrected models, respectively, and M was a baseline multi-trait model. Models have FE defined using NLEE of corresponding name.

**Figure S21:**
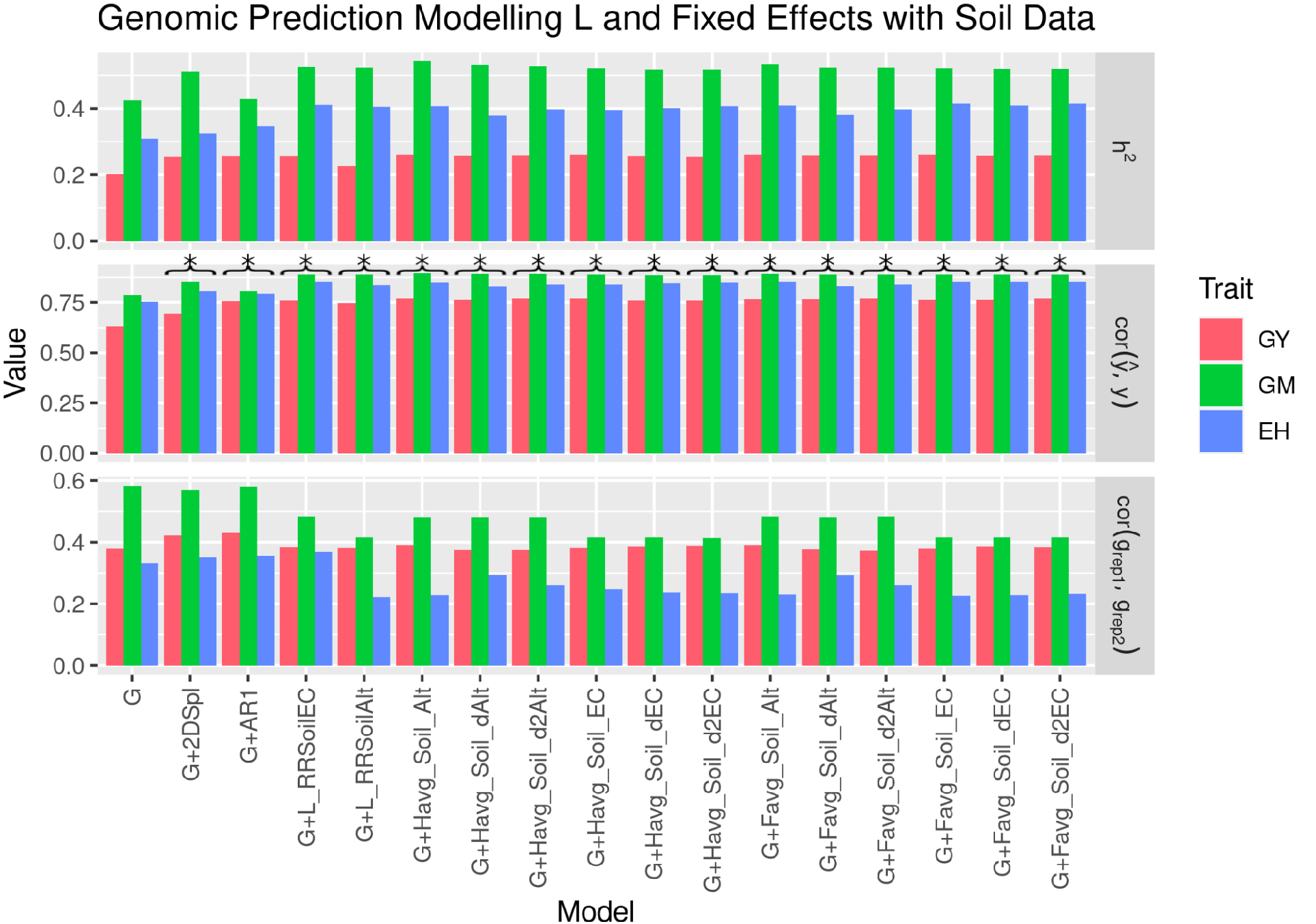
Genomic heritability (*h*^2^), model fit (cor(*ŷ,y*)), and genotypic effect estimation across replicates (cor(*g*_rep1_, *g*_Rep2_)) in 2019_NYH2 for GY, GM, and EH, with NLEE implemented as L or as FE using soil data.

**Figure S22:**
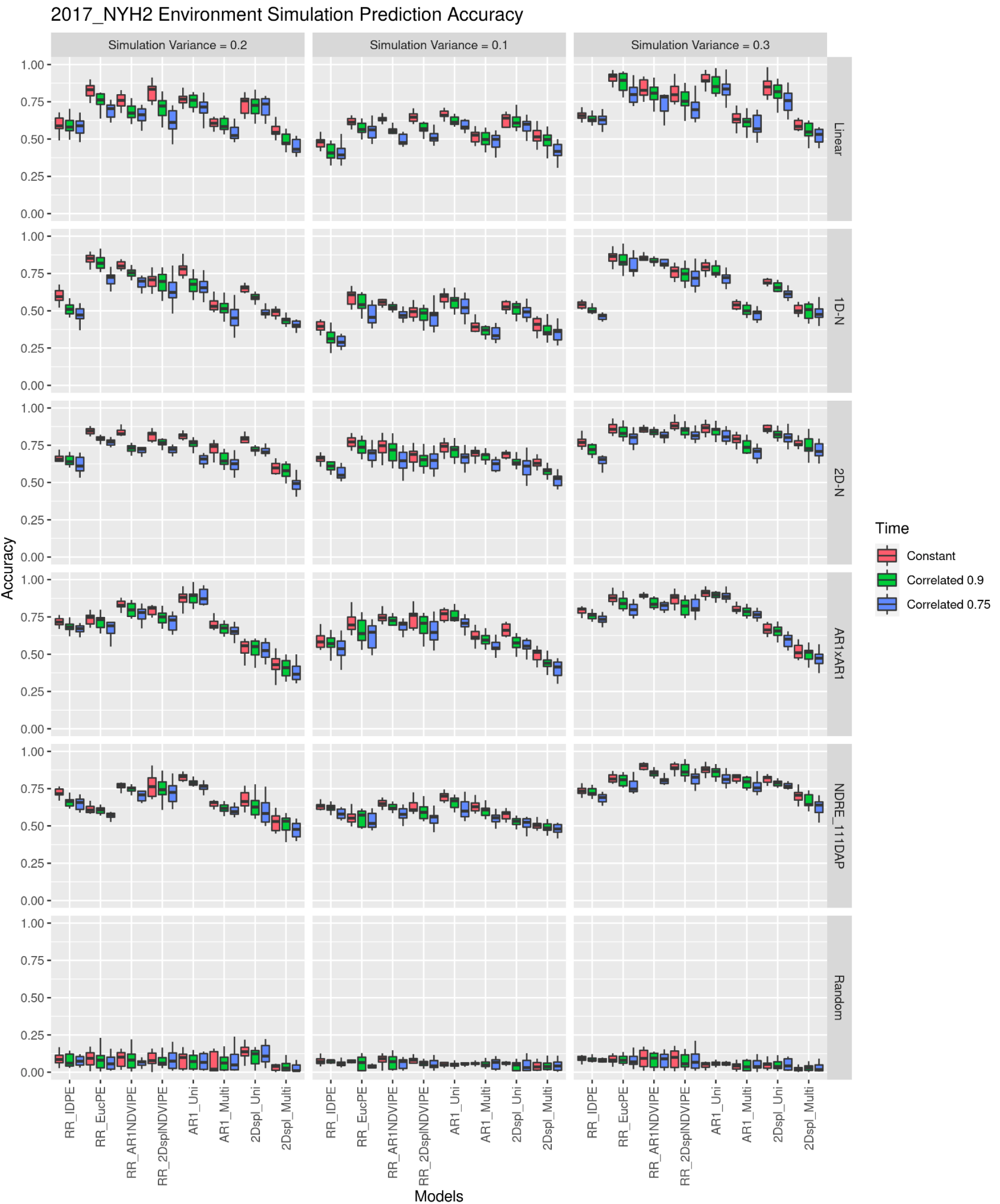
Prediction accuracy for six simulation processes (linear, 1D-N, 2D-N, AR1xAR1, random, and RD) were run ten times using the 2017_NYH2 NDVI values under a simulated environmental variance of 10%, 20%, and 30% and a simulated environmental effect that was 75%, 90%, and 100% correlated throughout the growing season. The RD scenario is illustrated as EC in this case. Prediction accuracy is the correlation of the simulated environmental effect and the model’s recovered effect.

**Figure S23:**
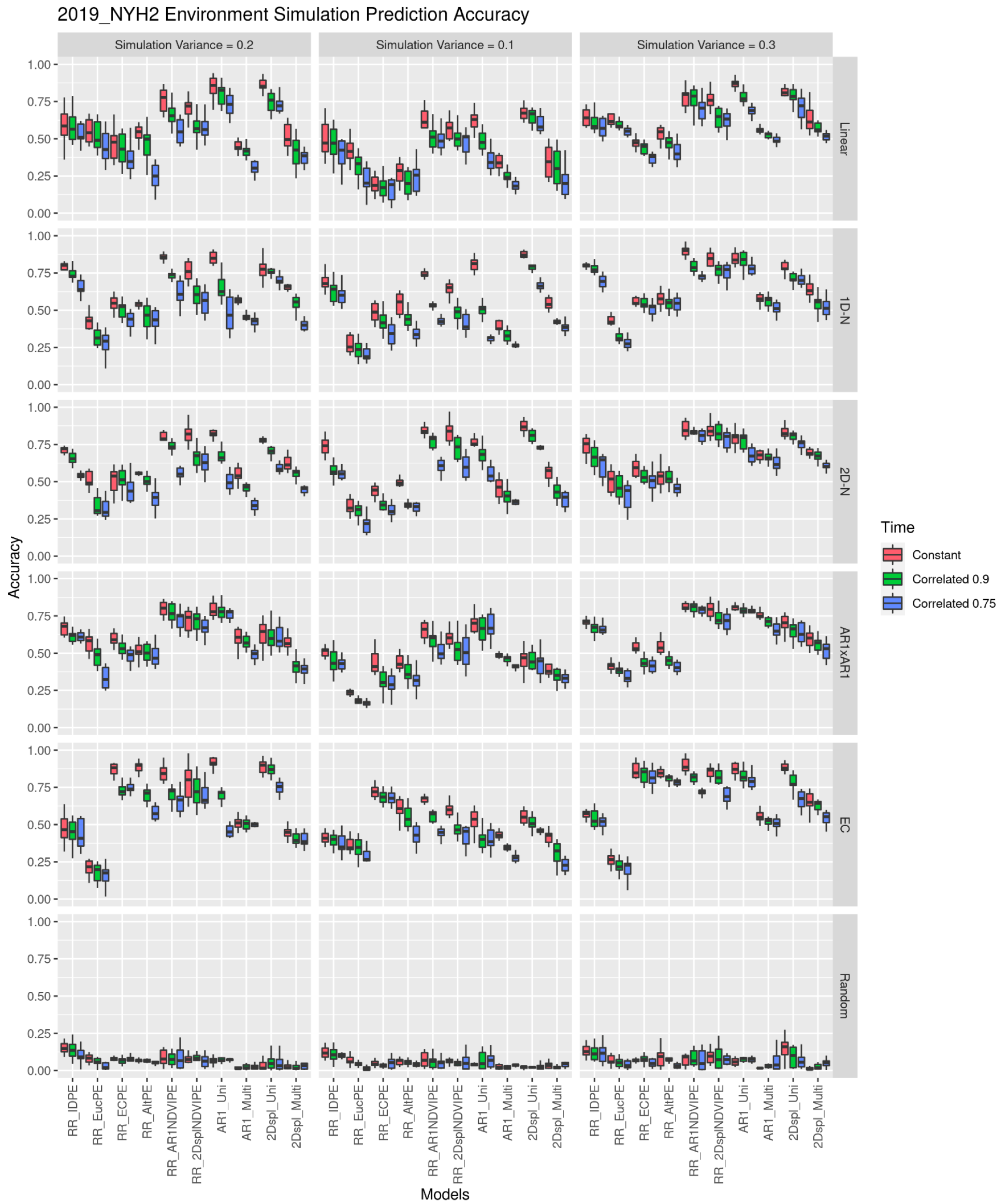
Prediction accuracy for six simulation processes (linear, 1D-N, 2D-N, AR1xAR1, random, and RD) were run ten times using the 2019_NYH2 NDVI values under a simulated environmental variance of 10%, 20%, and 30% and a simulated environmental effect that was 75%, 90%, and 100% correlated throughout the growing season. The RD scenario is illustrated as EC in this case. Prediction accuracy is the correlation of the simulated environmental effect and the model’s recovered effect.

**Figure S24:**
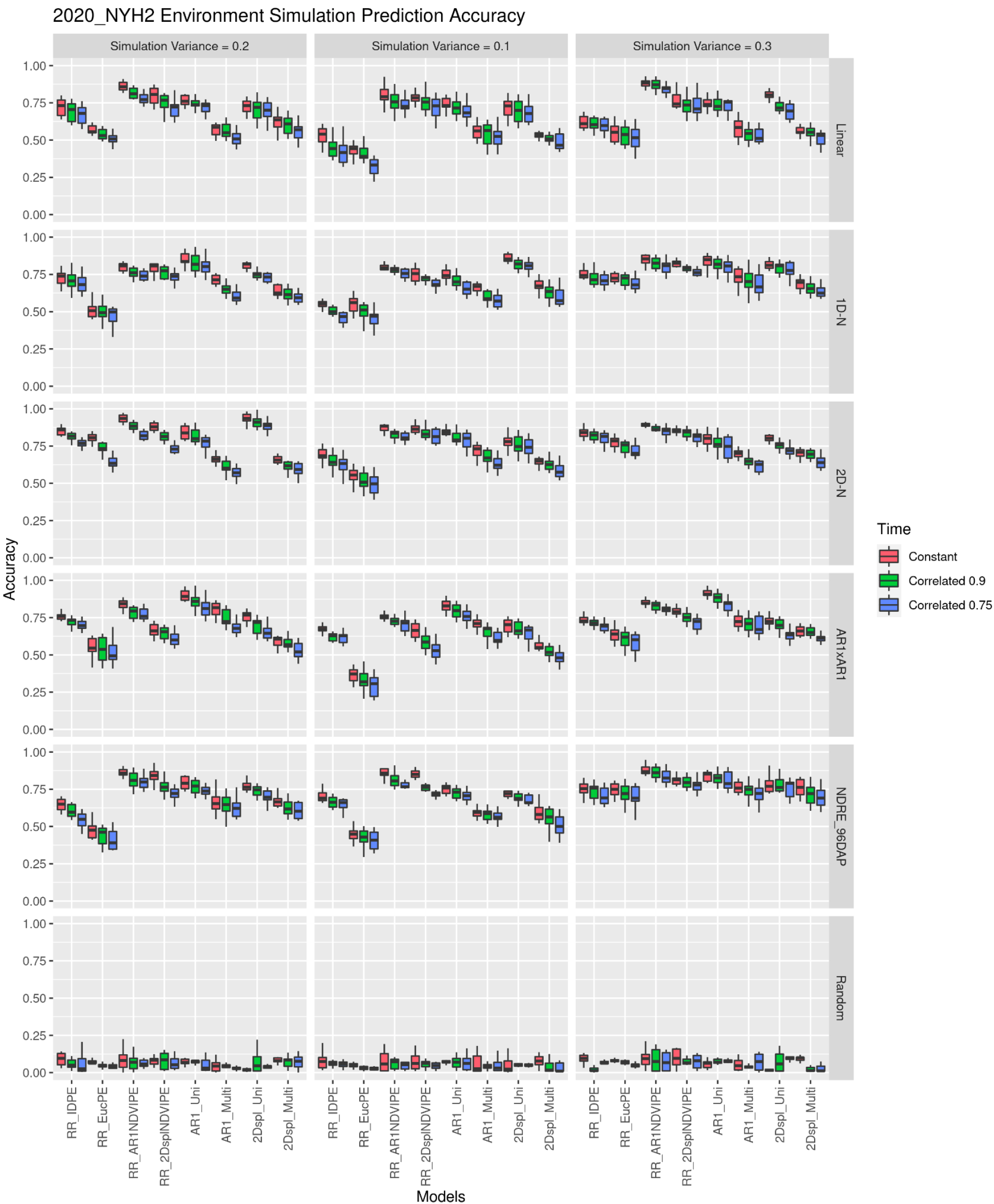
Prediction accuracy for six simulation processes (linear, 1D-N, 2D-N, AR1xAR1, random, and RD) were run ten times using the 2020_NYH2 NDVI values under a simulated environmental variance of 10%, 20%, and 30% and a simulated environmental effect that was 75%, 90%, and 100% correlated throughout the growing season. The RD scenario is illustrated as EC in this case. Prediction accuracy is the correlation of the simulated environmental effect and the model’s recovered effect.

